# Differentiating specific and non-specific protein-metabolite interactions using gradient open port probe electrospray ionization mass spectrometry

**DOI:** 10.1101/2024.03.07.583904

**Authors:** Xiaobo Tian, Gérard Hopfgartner

## Abstract

Native electrospray ionization mass spectrometry (ESI-MS) using nano ESI or desorption electrospray ionization (DESI) has been widely used to study interactions between macromolecules and ligands, usually protein-metabolite interactions (PMIs). In MS spectra the charge state distributions (CSD) of proteins differ between native and non-native conditions and based on this, we report a method that can differentiate specific protein-metabolite interactions from non-specific binding. Our approach is based on a 3D-printed open port probe electrospray system (gOPP-ESI) using mobile phase gradients (aqueous to methanol) for the ionization of protein/protein-metabolite complex. Notably, we found that a true protein-metabolite complex is more resistant to the denaturing effect of methanol compared to the free protein. This is corroborated by the observation that forming high charge states of protein-metabolite complexes requires higher proportions of methanol than free protein while, for non-specific complexes, there is no obvious difference in the CSD. Therefore, by comparing the changes in the CSD of free protein and protein-metabolite complex versus the increase of methanol, we can distinguish metabolites that specifically interact with the target protein. The approach is evaluated with well-characterized protein-ligand pairs, and we confirmed that cytidine phosphates, *N, N′, N″-*triacetylchitotriose, and fluvastatin are specific ligands for ribonuclease A, lysozyme, and beta-lactoglobulin respectively. However, cytidine-5’-triphosphate (CTP) interacts non-specifically with lysozyme and beta-lactoglobulin. We believe that after first-round native-MS assays to identify which metabolites cause mass shifts to the free protein, the gOPP-ESI-MS could be used as a quick second-round check to exclude non-specific binding and discover metabolites truly interacting with the protein of interest, reducing the number of candidates for subsequent validation experiments.

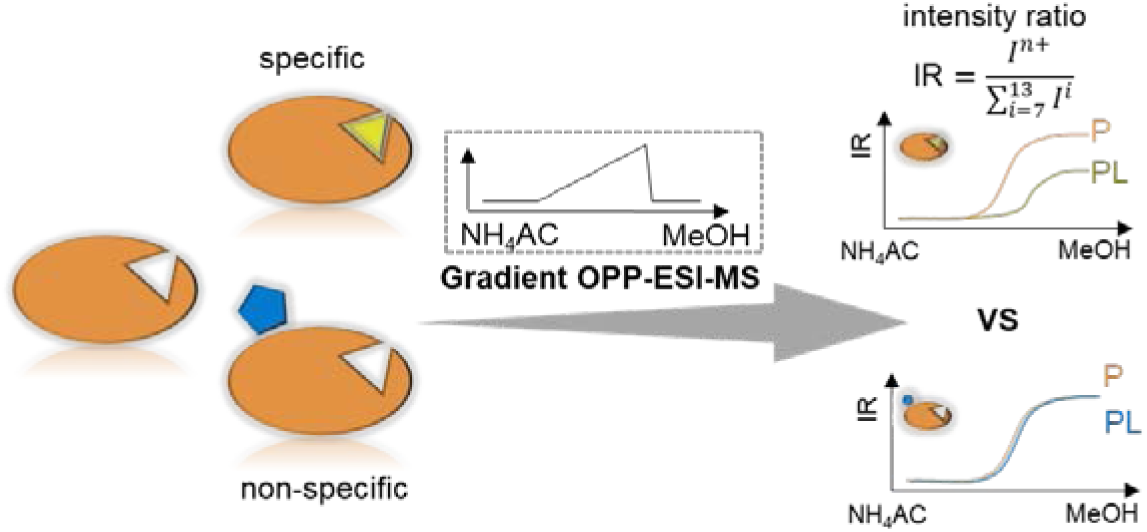

## INTRODUCTION

It is well known that most small molecules, like metabolites or drugs, produce effects through interactions with biomacromolecules, usually proteins. The investigation of protein-metabolite interactions (PMIs) is essential for deep understanding of the biological mechanism and has promise in drug discovery for screening hits for the target protein.^1^ A variety of techniques have been reported to study PMIs^2^, including surface plasmon resonance spectroscopy, isothermal titration calorimetry, nuclear magnetic resonance spectroscopy, and electrospray ionization mass spectrometry (ESI-MS).^3–6^ Unlike other methods, mass spectrometry has valuable advantages such as high sensitivity, high-speed analysis, low sample consumption, little-to-no sample preparation, etc., which make it a routine tool for protein analysis. Methods for studying PMIs based on MS can be roughly classified into two categories: metabolite-centric and protein-centric approaches.^7^

Metabolite-centric approaches use a known metabolite and look for protein targets^8^ and the most remarkable methods are those that do not require derivatization of the small molecules and directly analyze changes induced by PMIs with proteomics tools.^4, 8^ A common experimental routine is to use two groups, with and without ligands, and compare the proteomics data of these groups to determine ligand-induced changes when perturbations are introduced, such as heating^9^ and chemical denaturant.^10–12^ There have been many reports in this area, for instance, thermal proteome profiling^9, 13^ (TPP), chemical denaturation and protein precipitation^10^ (CPP), and solvent-induced protein precipitation^12^ (SIP), all of which seek PMIs based on alterations in protein stability caused by heating or chemical denaturants in the presence or absence of ligands. Limited proteolysis-coupled mass spectrometry^14, 15^ (LiP-MS) and pulse proteolysis^16^ enable the detection of - PMIs on the basis of the ligand binding-altered protease susceptibility of proteins. The stability of proteins from rates of oxidation^17, 18^ (SPROX), histidine-hydrogen deuterium exchange^11^ (His-HDX), and target-responsive accessibility profiling^6, 19^ (TRAP) explore PMIs relying on the difference in the accessibility of the oxidates/reactants to specific amino acids, like methionine, histidine, and lysine. Alternatively, protein-centric approaches start from a known protein, and search for interacting metabolites. One of the most widely used methods is the native ESI-MS,^20^ in which a metabolite is mixed with the target protein in a native-like solution, usually ammonium acetate (NH_4_Ac), and the mixture directly introduced to ESI-MS without separation/purification and while maintaining the PMIs during ionization. Subsequently, PMIs are explicitly detected by the mass shift of a protein-metabolite complex with respect to the free protein. In addition, quantifying the fractions of metabolite-interacted protein and free protein, information on stoichiometry and dissociation constants^21–23^ can also be inferred. With the simplicity of sample preparation, low sample consumption, high sensitivity, and high throughput, native ESI-MS has received a lot of attention in drug screening.^1, 2, 24, 25^ To optimize the native ESI-MS to explore PMIs, a variety of studies have been reported, including the type of ion source,^21, 26, 27^ the size of ESI tips,^28–31^ the buffer maintaining the native conditions,^32–34^ and solvent additives.^22, 35, 36^ However, only a few studies have dealt with the issue of non-specific binding between the target protein and the small molecules tested.^27, 37, 38^

A widely accepted ionization mechanism in native ESI-MS is the charged residue model^39–41^ (CRM) that assumes the ESI droplets shrink with solvent evaporation until all solvent molecules are gone and ions are generated. In this process the concentration of ligands and protein continues to increase which results in a higher likelihood of non-specific binding between the target protein and ligands compared to physiological conditions.^32, 37, 38^ Non-specific binding is structure independent and usually occurs on the solvent-exposed protein surface, which is consistent with the observed increase with the ligand concentration.^27, 30^ Thus, it might be more problematic with weak PMIs, because an molar excess of ligands is usually required to generate detectable amounts of PMIs, which promotes the occurrence of the non-specific binding. Klassen and coworkers proposed the reference protein method,^37, 38^ where a specific protein that does not interact with the tested ligands is used as an internal reference, and the fractions of ligand-adhered reference protein are used to correct the true PMIs. Ye and coworkers^27^ reported differentiating true PMIs from non-specific binding by acetonitrile-induced changes of the dissociation constant (Kd). For true PMIs, the Kd should decrease with an increasing proportion of acetonitrile in ESI, and vice versa for the non-specific binding.

The open port probe (OPP) was proposed by Van Berkel et al.^42^ and its application has been extended in various area.^43–45^ The core structure of the OPP is two coaxial tubes, the outer one receives solvents from pumps and the inner one aspirates samples into the electrospray ion source.^42, 46^ Samples can be independently introduced by pipette, syringe pump, or autosampler onto the dome of the OPP and are instantly aspirated into the ion source. To simplify OPP integration with different instruments (i.e. compatible with general ESI ion sources) and reduce costs (∼3 euros per piece of OPP), we produced devices with 3D printers and used them to qualitatively and/or quantitatively study metabolomics samples, thus eliminating carryover but retaining high speed sample introduction.^45, 46^

In the present study, we report a method that can differentiate true PMIs from the non-specific binding that is usually observed in native ESI-MS analysis. The method exploits the advantage of the OPP that the solvent can easily and continuously changed from native (NH_4_Ac solution) to denaturing (90% MeOH solution) conditions for ionization without disturbing sample introduction. With this approach we observed the important phenomenon that the stabilization induced by the PMIs makes the protein-metabolite complex have reduced intensity of high charge state ions compared to the free protein, which we then developed as method to identify true PMIs. To show proof-of-concept, we investigated well-studied protein-ligand pairs, including ribonuclease A, lysozyme and beta-lactoglobulin and the corresponding ligands cytidine phosphates, N, N′, N″-triacetylchitotriose and fluvastatin. This study revealed that, in contrast to true PMIs, non-specific binding between the target protein and ligands do not change the charge state distribution (CSD) compared to the free protein. Our approach may serve as a quick second-round check to exclude non-specific binding after the first-round native-MS assays, thus reducing the number of candidates to the target protein.

## EXPERIMENT SECTION

### Chemicals and Materials

Ribonuclease A (RNase A), lysozyme and beta-lactoglobulin, *N, N′, N″*-triacetylchitotriose (NTAC), cytidine-5’-triphosphate (CTP), cytidine-5’-diphosphate (CDP), cytidine-5’-monophosphate (CMP), cytidine, fluvastatin, formic acid, and ammonium acetate (NH_4_Ac) were purchased from Sigma-Aldrich. H_2_O, methanol (MeOH) and acetonitrile (ACN) for LC-MS were purchased from Chemsolute.

### Studies of protein-metabolite interaction under native or nonnative conditions

RNase A, lysozyme, or beta-lactoglobulin were dissolved in 25 mM NH_4_Ac to a concentration of 100 µM and the corresponding ligands, cytidine phosphates, NTAC, or fluvastatin added to a concentration of 1 mM. The mixtures were shaken at room temperature for 5 minutes before analysis and infused, with a syringe pump, onto the OPP at a rate of 3 µL per min. For the native condition, the OPP received the same solution of 10 mM NH_4_Ac from the LC pump. For the nonnative condition, the OPP received the solutions via a LC gradient: 0-5 min, constant at 100% of 10 mM NH4Ac; 5-10 min, change from 100% of 10 mM NH_4_Ac to 100% of H_2_O containing 0.1% formic acid; 10-20 min, change from 0 to 90% of MeOH. 20-25 min, equilibrate at 100% of 10 mM NH_4_Ac.

### OPP-ESI-MS

MS analyses were performed on a TripleTOF 5600 (Sciex, Concord, ON) integrated with a 3D-printed OPP. The outer tube of the OPP was connected to an LC pump (Shimadzu LC-30AD) and the inner tube was connected to the Turbo V ion source. Experiments were conducted in positive ion mode with a TOF-MS experiment (*m/z* 800-2500). Ionization settings were as follows: GS1 = 80; GS2 = 45; curtain gas = 30; TEM = 300 °C and ISVF = 5000 V. Data were acquired with Analyst software (version 1.6.2).

### Data analysis

The intensity ratio (IR) for a single charge state in a specific spectrum was calculated by dividing its intensity by the summed intensity of all charge states, IR = I^n+^/I^total^. The intensity ratio of for each charge was plotted versus the gradient. The binding ratio (BR) was determined by dividing the total intensity (all charge states) of protein-metabolite by the sum of the total intensity of protein-metabolite and the total intensity of unbound-protein, BR = [PL]/([PL] + [P]). For the Kd evaluation of CTP and CDP to RNase A, the ratio, *R*, of the ligand-bound protein to the free protein was calculated based on intact masses in the deconvoluted spectra that were reconstructed by the embedded Bio Tool Kit in PeakView Software (version 2.2).

## RESULTS AND DISCUSSION

### Conception of the gradient-OPP-ESI-MS approach

True interactions between proteins and metabolites usually tighten/lock the protein structure and are the basis for metabolite-centric approaches that identify bound proteins by comparing protein stability with and without the target metabolite.^11^ This difference is even more pronounced in the presence of perturbations such as heating and chemical denaturants. Meanwhile, CSD is widely used to study protein structure changes since an unfolded protein usually has more high charge state ions than its folded forms.^47–49^ Thus we reasoned that involving denaturing perturbations in native ESI-MS could facilitate the identification of metabolites truly interacting with the target protein. More specifically, we hypothesized that if the proportion of MeOH in ESI was increased, the protein-metabolite complex would show a more significant CSD difference compared to the free protein. In contrast, non-specific interactions are usually on the surface of proteins and should have less influence on structure, and hence less effect on the CSD. Here, we describe the gOPP-ESI-MS approach which differs from conventional native ESI-MS because the OPP allows ESI solvent to be changed while keeping the protein/protein-metabolite complex under native conditions. As shown in **Figure 1**, the protein with and without ligand is introduced onto the OPP in a dropwise way and instantly aspirated into the ESI source. Increasing the proportion of MeOH in the OPP solvent with a gradient gradually increases the denaturing effect of the solvent. As a result, proteins will be unfolded stepwise and the CSD of the protein will be broadened and shift to higher charge states. However, due to the stabilization brought by the PMIs, the protein-metabolite complex is expected to be more resistant or show lower intensities in high charge states than that of the free protein. Although increasing MeOH could break some PMIs, the intensities of the protein-metabolite ions do not necessarily decrease since the ESI response sensitivity increases with higher proportions of MeOH.^50, 51^ Remarkably, our approach to screening PMIs based on the distribution of charge states is not affected by peak intensities.

**Figure 1.**
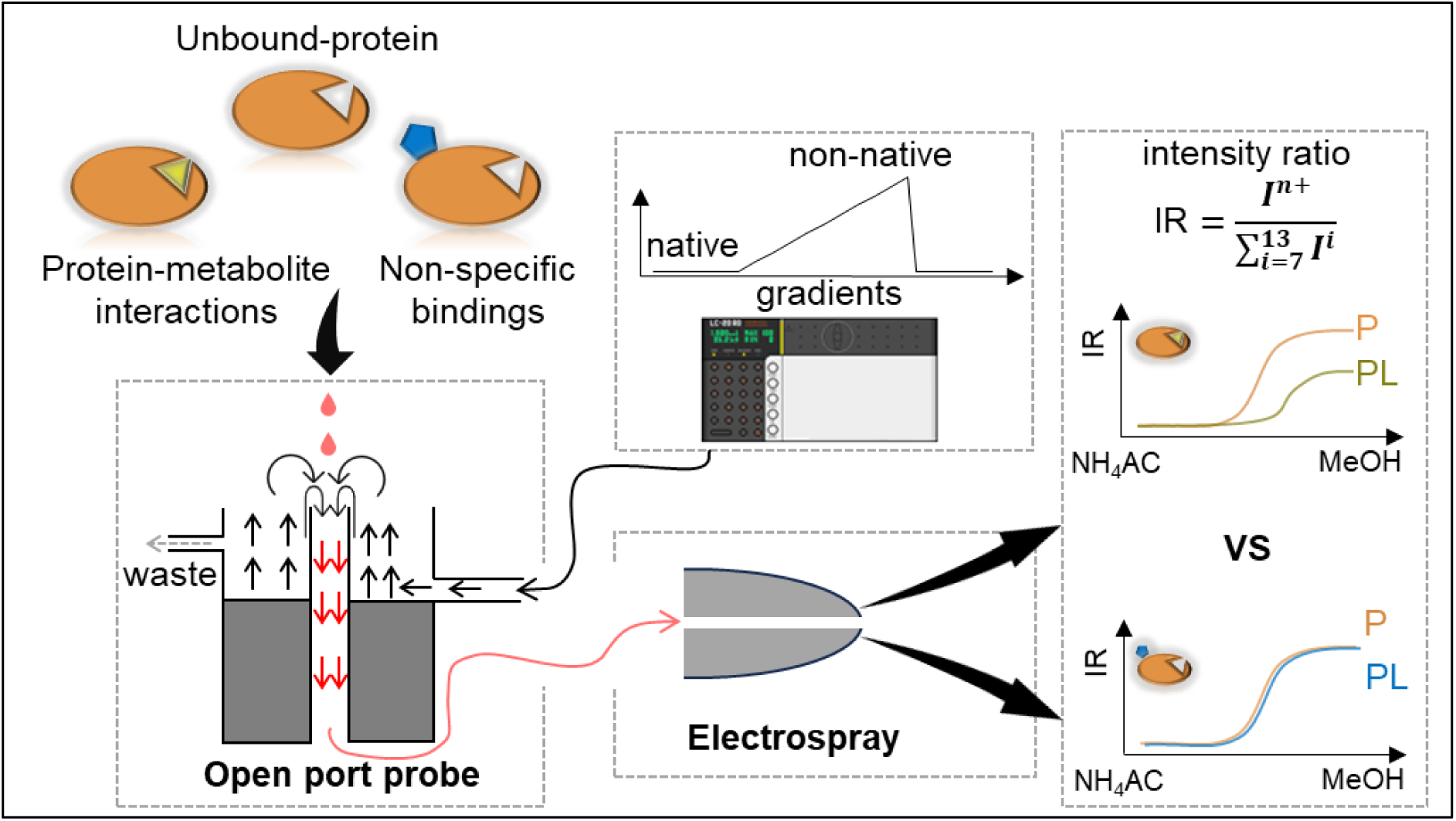
Schematic showing the gOPP-ESI-MS approach for identifying metabolites interacting with the target protein. The system includes four parts: an LC pump, the sample introduction, the OPP, and an ESI-QqTOF MS. As depicted, protein samples with and without ligands are introduced by pipette or syringe pump to the OPP where solutions with a constant composition or gradient changes are delivered by an LC pump. Under different denaturing strength, the protein and protein-metabolite complex show differences in CSD, especially in higher charge states.

### Probing protein-metabolite interaction under constant native conditions

As a variant of native ESI-MS, we started by testing the performance of the gOPP-ESI-MS approach to measure protein metabolite complexes under the native condition of 10 mM NH_4_Ac. We selected the well-characterized proteins^21, 27^ RNase A, lysozyme and beta-lactoglobulin. For RNase A three positive ligands (CTP, CDP, and CMP) and a negative ligand (cytidine) were separately incubated with RNase A and infused onto the OPP by a syringe pump. As shown in **Figure 2 A-D**, both the free RNase A and three complexes of RNase A with cytidine phosphates exhibited a narrow CSD centered on charges of 7+ and 8+, which indicates that they are in the folded status. To simplify the *R* calculation, the ratio of the ligand-bound protein to the free protein, the deconvolution was conducted, and the *R* is calculated based on intact masses. For CTP, CDP, and CMP, the *R* values are 1.48, 0.60, and 0.31 respectively, which is consistent with the affinity orders reported previously.^21^ No complex was observed for cytidine.^52^ The affinity orders were further investigated in competitive and noncompetitive experiments, see in **Figure S1**, the measured orders are consistent with each other. Furthermore, as shown in **Figure S2**, the Kd of CTP and CDP were measured in titration assays according to the method reported by Daniel et al.^22, 53^, in which the concentration of RNase A was constant (5 µM) and the ligand investigated at a series of concentrations from low to high (0.5 – 50 µM). The measured Kd for CTP and CDP were 10.9 and 38.61 respectively, which is slightly higher than the value reported in the earlier study.^21^ This may be due to the temperature of 300 degrees required for sufficient and stable ionization in our ion source, while this temperature can be decreased below 100 degrees in nano ESI. Note, however, that the focus of the present work is not determining Kd but rather the capability to distinguish between true PMIs and non-specific binding.

To show the issues of non-specific binding, we selected a true-positive ligand (NTAC) and a false-positive ligand (CTP) for lysozyme.^27^ In **Figures 2E** and **F**, the free lysozyme and the complexes with NTAC and CTP are centered on the charge of 8+. The *R* value of NTAC is 0.59 which denotes a relatively strong binding to lysozyme but the false-positive ligand, CTP, also shows a low but reliable peak at the *m/z* of lysozyme-CTP with a *R* value around 0.05 (**Figure 2F**). Moreover, In **Figures 2G** and **H**, we also investigate the non-specific binding of CTP and beta-lactoglobulin and compare it with fluvastatin which was reported to specifically bind to beta-lactoglobulin.^54^ Like these, just based on the mass shifts, we could not exclude CTP as the real ligand for lysozyme and beta-lactoglobulin.

**Figure 2.**
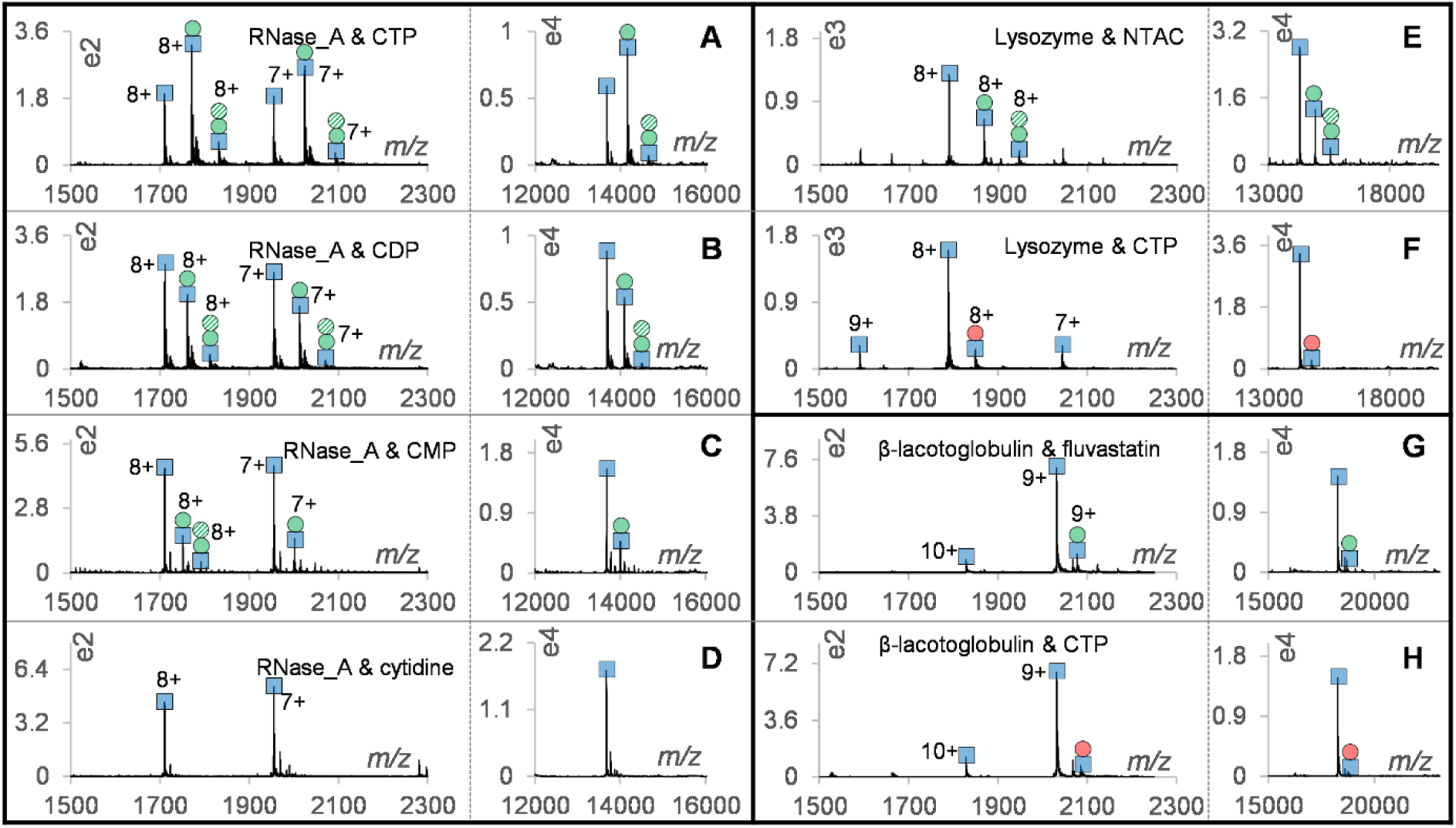
Probing PMIs under the constant native condition of 10 mM NH_4_Ac. In each panel, the left part is the raw spectrum, and the right part is the corresponding reconstructed spectrum after deconvolution based on multiple charge states. The proteins and ligands are freshly mixed and incubated for 5 minutes, then infused onto OPP by a syringe. The investigations of RNase A (100 µM): **A** with CTP (1 mM), **B** with CDP (1 mM), **C** with CMP (1 mM), and **D** with (1 mM) cytidine. The investigations of lysozyme (100 µM): **E** with NTAC (1 mM) and **F** with CTP (1 mM). The investigations of beta-lactoglobulin (50 µM): **G** with fluvastatin (500 µM) and **H** with CTP (500 µM). The blue square, green solid circle, green dashed circle, and red solid circle indicate the protein, the true-positive ligand, the second binding and the false-positive ligands respectively.

### Charge state distribution changes with increasing methanol in the OPP solution

The core finding in this work is that we can differentiate true PMIs from non-specific binding by comparing the changes in the CSD of the free protein and protein-metabolite complex versus the increasing amount of methanol. To show the capability of modulating CSD from native to unfolded status, protein solutions of RNase A and myoglobin were continuously infused onto the OPP while the proportion of MeOH in the OPP solution was gradually increased from 0 to 90% with an LC pump. Overall, the peak intensities of proteins increased more than five times over the gradient, as shown by the base peak chromatographs (BPCs) in **Figures 3A** and **S3**. For RNase A, **Figure 4A**, as mentioned previously, the CSD of native RNase A was centred on charges of 7+ and 8+ with 0% MeOH (100% 10 mM NH_4_Ac). Charges of 9+ and 10+ appeared around 5% MeOH (**Figure 4B**), followed by the higher charge states (10+ to 14+) which increased with higher MeOH (**Figures 4C** and **D**). To show the CSD changes induced by ligand binding, the complex of RNase A with CTP was investigated at the same conditions (**Figures 4E** to **H**). In 10 mM NH_4_Ac, the charge states didn’t change and centred on charges of 7+ and 8+ too (**Figure 4E**). While with the higher proportion of MeOH, the complex of RNase A with CTP showed higher IR for charges 10+ and 9+ (**Figure 4G**). An interesting phenomenon is that, with higher MeOH, we observed phosphate adducts on RNase A. This phenomenon was reported in a previous work.^55^ Further, we investigated another model protein, myoglobin, that contains a ligand, heme in the native state.^27^ As shown in **Figure 3B**, with 10 mM NH_4_Ac the CSD of the heme-bound myoglobin centered on charges 8+ and 9+, meanwhile, almost no heme was observed. With the solvent changed to 100% H_2_O containing 0.1% formic acid, **Figure 3C**, the heme-bound myoglobin and the free myoglobin exist simultaneously in the spectrum, and we estimated that more than 50% of complexes are already dissociated. However, the free heme did not obviously increase (**Figure 3B** vs **C** in right parts) and we speculate this is because the OPP 100% aqueous solution is not appropriate for the ionization of heme. In contrast, with higher levels of MeOH (**Figures 3D** and **E**), the intensity of heme increased at least 10 times (**Figure 3B** vs **E** in right parts) which is much higher than the intensity changes of the free myoglobin. The appearance of a broad CSD and high charge states reveal the unfolding of the protein that makes more ionization sites accessible and results in a gradual increase in spectrum intensity.

**Figure 3.**
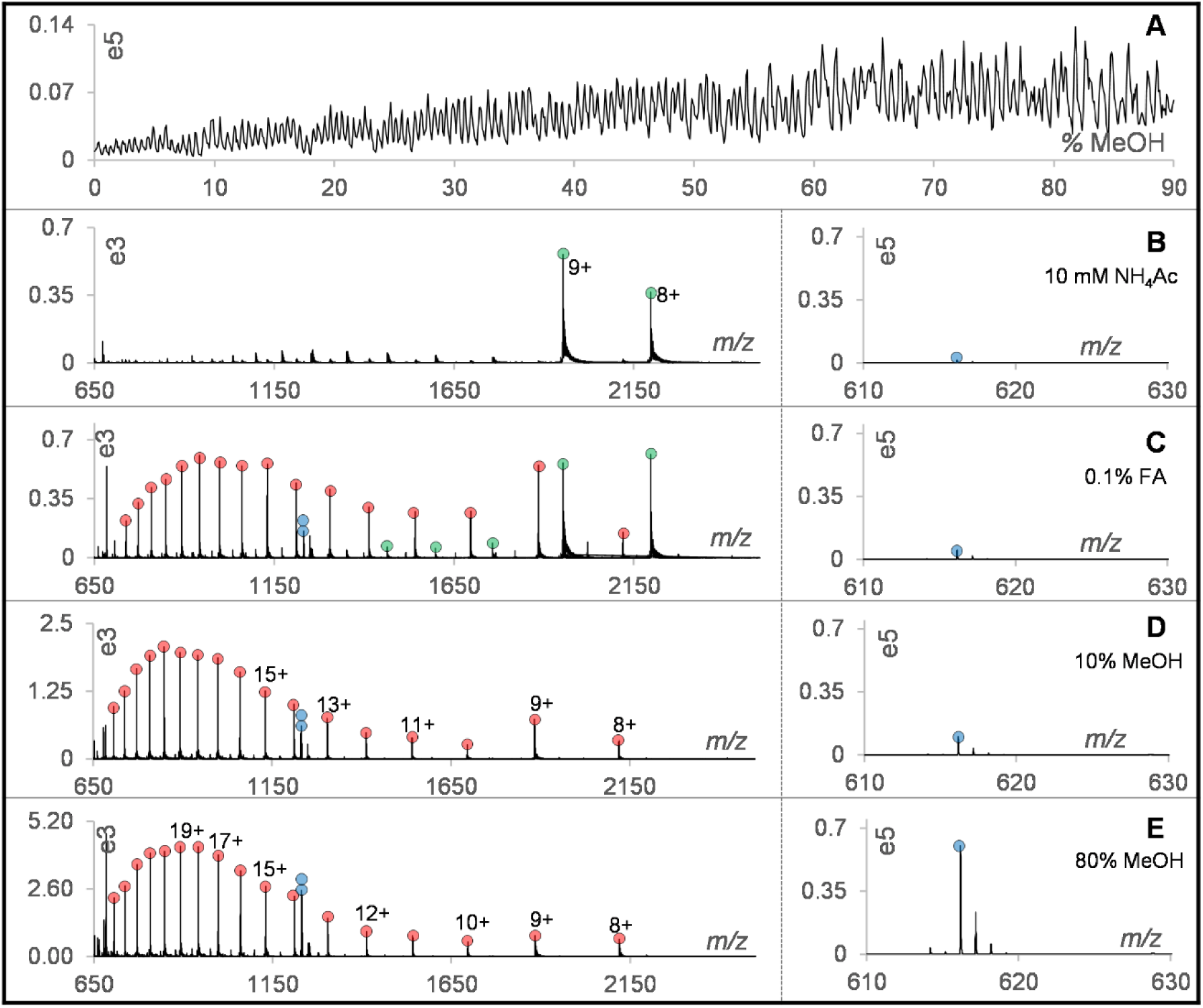
Modulating CSD of myoglobin by increasing the proportion of methanol in the OPP solvent. **A**. the BPC of the heme-bound myoglobin and/or the free myoglobin over the gradient from 0 to 90% MeOH. Raw spectra of the heme-bound myoglobin and/or the free myoglobin at various proportions of MeOH: **B**. 100% 10 mM NH_4_Ac, **C**. 100% H_2_O containing 0.1% formic acid, **D**. 10% MeOH, **E**. 80% MeOH. The green and red circles indicate the CSD of the heme-bound myoglobin and the free myoglobin respectively. The blue circle represents the peak of heme (*m/z* of 616.17) and the “double-blue circle” denotes the peak of (2M+H)^+^ at *m/z* 1231.27. In panels **B** to **E**, the left part is the spectrum of the free myoglobin or the heme-bound myoglobin and the right part shows the heme (*m/z* of 616.17) with a constant y scale.

**Figure 4.**
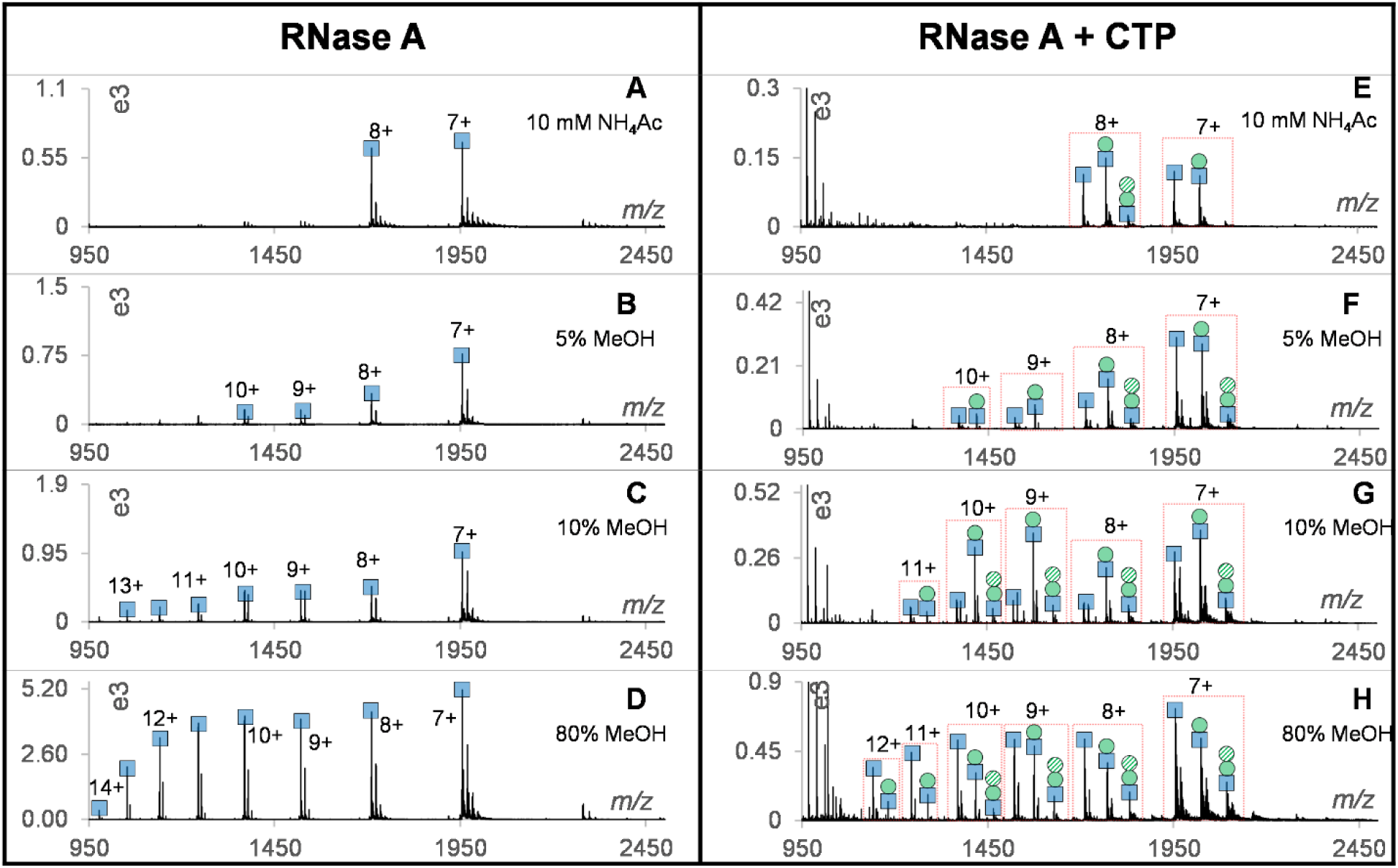
Modulating CSD of RNase A by increasing the proportion of methanol in the OPP solvent. Spectra of RNase A at various proportions of MeOH: **A**. 100% 10 mM NH_4_Ac, **B**. 5% MeOH, **C**. 10% MeOH, **D**. 80% MeOH. Spectra of CTP-bound RNase A and/or free RNase A at various proportions of MeOH: **E**. 100% 10 mM NH_4_Ac, **F**. 5% MeOH, **G**. 10% MeOH, **H**. 80% MeOH. The “double peak” in panels are adducts with phosphate (*m/z* shift of 98.3) and are most intense in panels **C** and **D**. The blue square, green solid circle and green dashed circle represent the protein, the first binding and the second binding respectively.

### Reproducibility evaluation and CSD differences between the free and CDP-bound RNase A

To evaluate the reproducibility of the gOPP-ESI-MS approach, triplicate analyses of the interaction between CDP and RNase A were investigated under the constant native condition (10 mM NH_4_Ac) and a gradient that ranges from native to non-native conditions. From **Figure S4**, we know around 38% of RNase A was bound with CDP in 10 mM NH_4_Ac and the ratios remain unchanged during the process, which implies the sample introduction by syringe and OPP-ESI ionization are reproducible enough for further analysis. In addition, the intensity ratio of all charge states remains unchanged under the native conditions (**Figure S5**). For the analyses under the gradient, as shown in **Figure 5A**, the binding ratio started to decrease around 8 minutes (at 40% 10 mM NH_4_Ac and 60% H_2_O containing 0.1% formic acid) and stayed constant at 15% after 12 minutes even with 90% MeOH. The dots related to the three replicates are close to each other, indicating that changing the OPP solution does not influence reproducibility.

Unlike the binding ratio, as shown in **Figure S6**, the intensity ratios of most charge states didn’t change obviously before 10 minutes, except the charge 8+ (**Figure 5E**) that roughly decreased from 50% to 20%. Besides, only the charges 7+, 9+ and 10+ slightly increased but no change for higher charge states 12+ to 14+. This suggests the need to introduce stronger perturbations to manifest the CSD difference. In line with that, after 10 minutes, with increasing the proportion of methanol in the OPP solution, high charge states 11+ to 14+ appeared gradually and reached the maximum around 17 minutes (**Figures 5B** to **E**). To illustrate the CSD difference caused by PMIs in details, in **Figure S6**, we named the RNase A in the control sample containing RNase A without CDP as “control-RNase A” and named the free RNase A in the experimental sample containing both RNase A and CDP as “unbound-RNase A”. As depicted in **Figure S6**, the CSD of control-RNase A and unbound-RNase A almost overlapped together in the gradient, which proves the excess free ligand in the experimental sample, CDP, does not affect the CSD of RNase A. Given this, we can confidently infer that changes in the CSD of the RNase A-CDP complex are due to their interactions, which may tighten/lock the structure of RNase A and alter the accessibility of ionization sites. As we expected, **Figures 5B** to **E**, the RNase A-CDP complex did show differences in CSD compared to unbound-RNase A. Especially for high charge states, 12+ and 13+, the distinction is more pronounced. They appear approximately 3 minutes later and have lower intensity ratios than the unbound-RNase A. The anti-denaturing effect of PMIs was exemplified in a specific spectrum, **Figure S7**, which shows the CSD difference between RNase A-CDP complex and unbound-RNase A. This finding is intriguing and may be useful in identifying metabolites that truly interact with target proteins.

**Figure 5.**
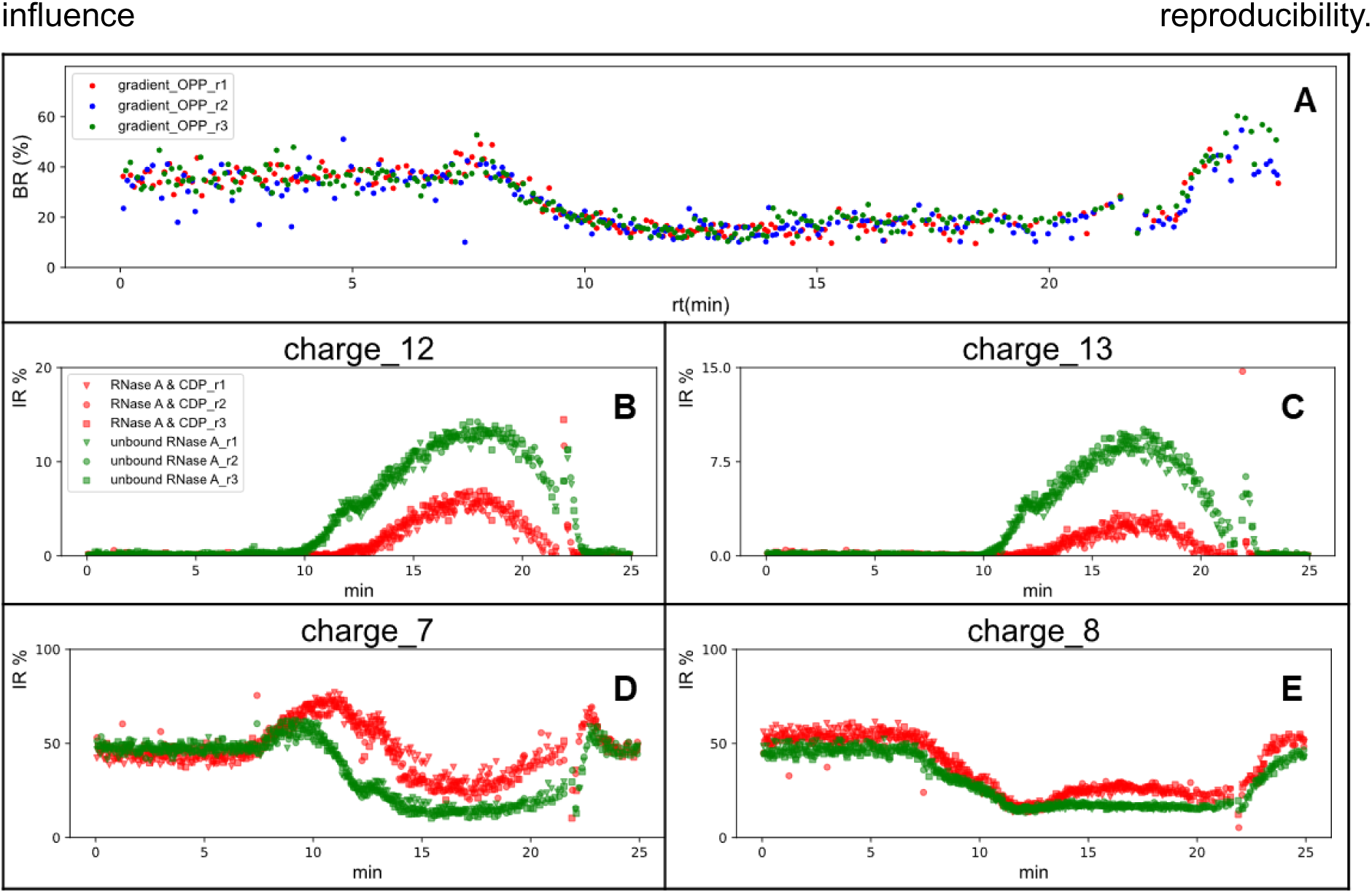
Reproducibility of the gradient-OPP-ESI-MS approach. **A**. triplicate analyses of CDP interacting with RNase A with a MeOH gradient. Binding ratio is the ratio [PL]/([PL] + [P]). Intensity ratios of specific charge states over the gradient: **B**. charge 12+, **C**. charge 13+, **D**. charge 7+ and **E**. charge 8+. Gradient: 0-5 min, constant at 100% 10 mM NH_4_Ac, 5-10 min, change from 100% 10 mM NH_4_Ac to 100% H_2_O containing 0.1% formic acid, 10-20 min, change from 0 to 90% MeOH. 20-25 min, equilibrate at 100% 10 mM NH_4_Ac. BR is binding ratio and IR is intensity ratio.

### Differentiating true PMIs and non-specific binding

To demonstrate the effectiveness of the gOPP-ESI-MS approach in differentiating between true PMIs and non-specific binding, we further studied lysozyme and beta-lactoglobulin with the corresponding true-positive and false-positive ligands. Consistent with previous reports,^27^ the true PMIs of lysozyme-NTAC and beta-lactoglobulin-fluvastatin showed a downward trend for binding ratios with higher proportions of methanol in the OPP solution (**Figures S8** and **S9**). In contrast, the non-specific binding of lysozyme-CTP and beta-lactoglobulin-CTP, were unchanged or even slightly increased. In addition, we noted that the true PMIs and non-specific binding have distinctive behaviour in CSD over the gradient, please see CSD of all charge states for lysozyme in **Figure S10** and for beta-lactoglobulin in **Figure S11**. For low charge states, like charge 9+ for lysozyme and charge 10+ for beta-lactoglobulin, the difference between PMIs and non-specific binding is not obvious (**Figures 6B** vs **G**). However, a gratifying discovery is that the intensity ratio differences are greatly pronounced for high charge states, such as 11+ and 12+ for lysozyme (**Figures 6D** vs **I** and **E** vs **J**) and charge 12+ and 17+ for beta-lactoglobulin (**Figures 7B** vs **G** and **E** vs **J**). Compared to the non-specific binding, high charge states of true PMIs arise at high proportions of MeOH in the OPP solution and are less intense. The findings are consistent with the hypothesis that binding to the corresponding true-positive ligand makes protein more resistant to unfolding, but non-specific binding, usually on the surface of proteins, should have less or no influence on the structure. Furthermore, in **Figure S11**, an interesting phenomenon was observed that for beta-lactoglobulin, the occurrence of charges 11+ to 13+ started from 10 minutes, but charges of 14+ to 18+ didn’t increase until 15 minutes. The two isolated uptrend appear to indicate it has two unfolding stages, which might be interesting for structural biology research.

**Figure 6.**
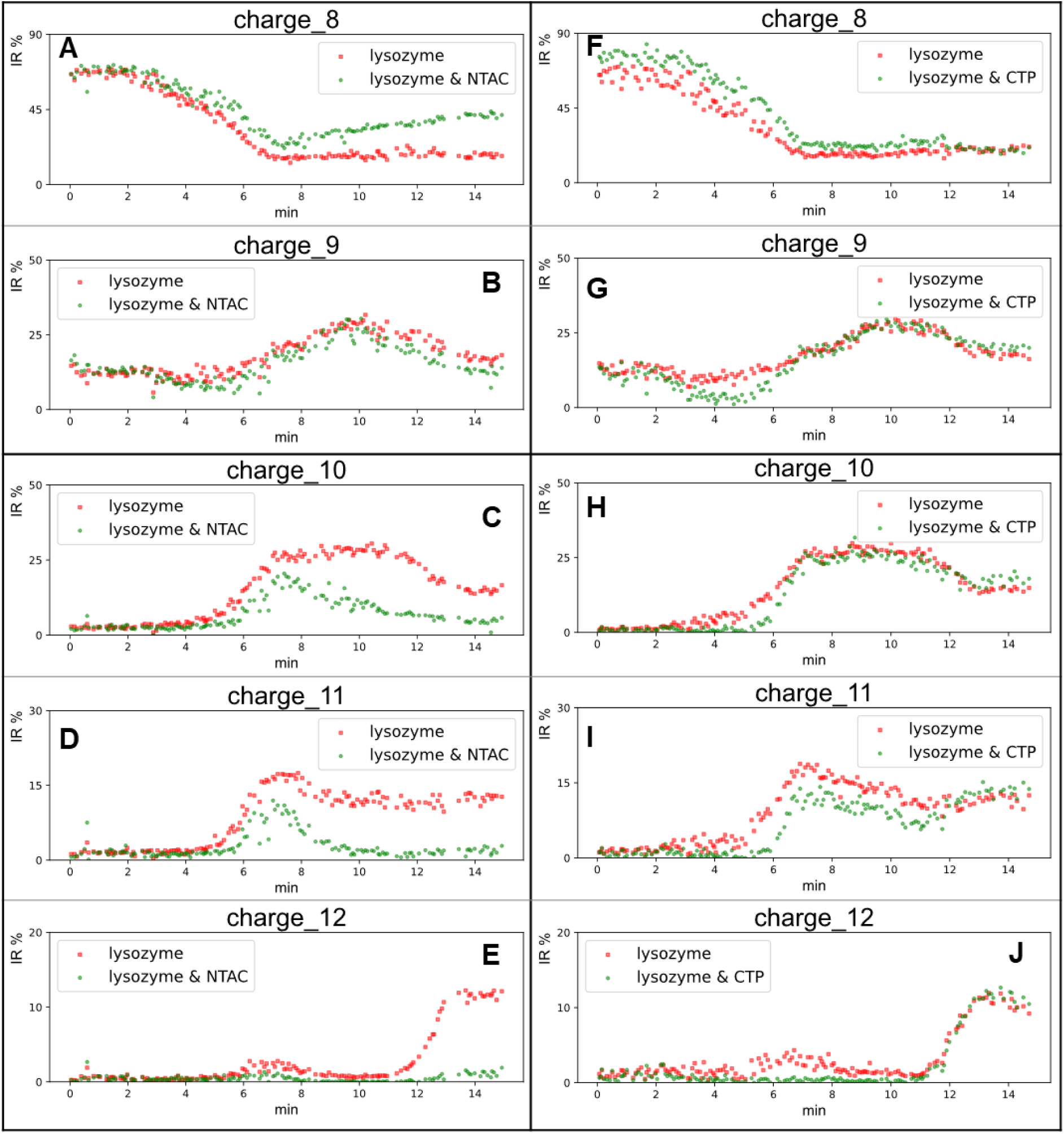
Differentiations between the true PMIs and non-specific bindings. Intensity ratio changes of lysozyme-NTAC in specific charge states: **A** to **E**. are charges 8+, 9+, 10+, 11+ and 12+; intensity ratio changes of lysozyme-CTP in specific charge states: **F**. to **J** are charges 8+, 9+, 10+, 11+ and 12.

**Figure 7.**
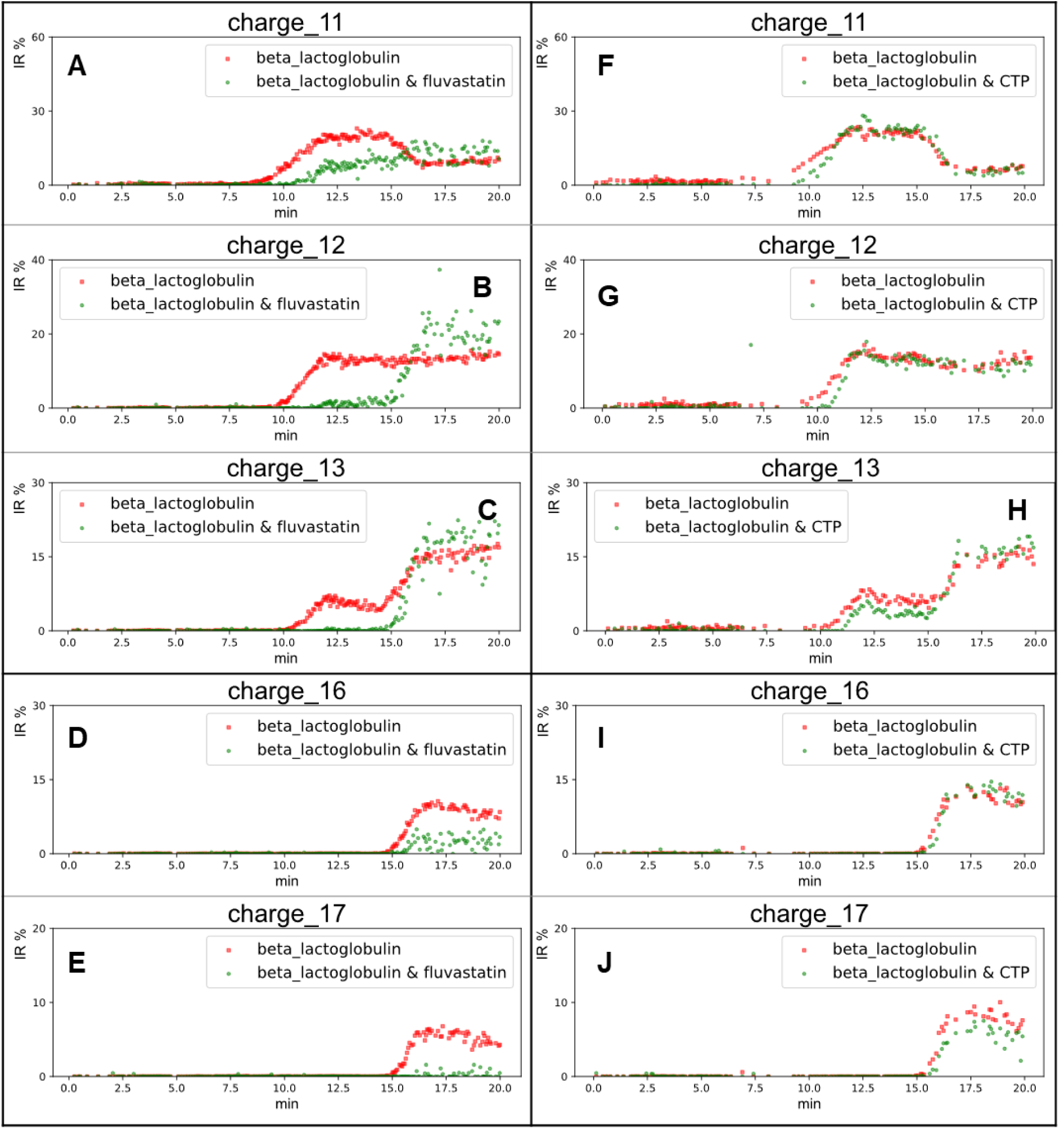
Differentiations between the true PMIs and non-specific bindings. Intensity ratio changes of beta-lactoglobulin and fluvastatin in specific charge states: **A** to **E**. are charges 11+, 12+, 13+, 16+ and 17+; intensity ratio changes of beta-lactoglobulin and CTP in specific charge states: **F**. to **J** are charges 11+, 12+, 13+, 16+ and 17+.

Conventional native ESI-MS approaches determine binding by mass shifts of the protein-metabolite complex with respect to the free protein. This method is widely used due to its simplicity and high throughput. However, non-specific binding that also cause mass shifts of the free protein cannot be excluded and a method that differentiates true PMIs and non-specific binding is therefore highly desirable. Reference protein methods^37, 38^ utilize a reference protein that does not interact with the tested ligands to correct the true PMIs, but requires the careful selection of reference proteins and may also introduce bias, since non-specific binding between different proteins and ligands vary from case to case. Involving perturbations, such as heating and chemical denaturants, to make the difference induced by PMIs more pronounced is common in metabolite-centric approaches, but rarely applied to native ESI-MS. This may be because of the belief that introducing perturbations would disrupt the PMIs or because of the lack of simple and effective devices to implement this approach. These needs are well matched with our 3D-printed OPP system that can easily and continuously change solvents from native to denaturing conditions with a programable LC pump. Since sample introduction of the protein/protein-metabolite complex is independent of the OPP, they can be kept under native conditions until the moment they are introduced, which in turn enables many selections of the OPP solutions. Notably, besides the proportion of organic solvents, the gOPP-ESI-MS can be easily extended to explore the influence of other perturbations on the PMIs, like pH, the proportion of detergent or concentration of buffer. Since PMIs are usually based on various forces, such as hydrogen bonds, hydrophobic interactions, electrostatic interactions, and Van der Waals forces, we anticipate that different perturbations may be used to study different types of interaction between protein and metabolites, for example acid/base for ionic interaction, urea for hydrogen bonding, and acetonitrile for hydrophobic interactions. Moreover, compared to the nano DESI that requires careful arrangement of the angles and alignments of the DESI probe, sample introduction capillary, and MS inlet capillary,^21^ the OPP simply mounts on a general ESI ion source with commonly used peek tubes and direct analysis of protein/protein-complexes can be achieved.

## CONCLUSIONS

Here, prompted by the desire to exclude non-specific protein-ligand binding, we propose to introduce perturbations, including organic solvents, chemical denaturants, acids, or bases, into the study PMIs by native ESI-MS. The approach was implemented with a 3D-printed OPP system coupled with ESI-MS, and we found that true PMIs and non-specific binding have distinctive CSD behaviour under conditions changing from native to non-native which we then developed as a criterion to screen the true PMIs. The differentiation between true-positive and false-positive ligands was exemplified with well-studied protein-ligand pairs, including RNase A, lysozyme and beta-lactoglobulin and the corresponding ligands. The gOPP-ESI-MS approach can be used as a quick second-round assay to exclude non-specific binding after conventional native ESI-MS experiments, and we believe that this approach holds the promise of avoiding false-positive ligands, thus further narrowing the range of drug candidates for the target proteins.

## Supporting information

Supplementary Infos

## ETHICAL STATEMENT

Does not apply.

## REFERENCES

(1) Gavriilidou, A. F. M.; Sokratous, K.; Yen, H.-Y.; De Colibus, L. High-Throughput Native Mass Spectrometry Screening in Drug Discovery. Frontiers in Molecular Biosciences 2022, 9, Review. DOI: 10.3389/fmolb.2022.837901.

(2) Woods, L. A.; Dolezal, O.; Ren, B.; Ryan, J. H.; Peat, T. S.; Poulsen, S.-A. Native State Mass Spectrometry, Surface Plasmon Resonance, and X-ray Crystallography Correlate Strongly as a Fragment Screening Combination. Journal of Medicinal Chemistry 2016, 59 (5), 2192–2204. DOI: 10.1021/acs.jmedchem.5b01940.

(3) Stincone, P.; Naimi, A.; Saviola, A. J.; Reher, R.; Petras, D. Decoding the molecular interplay in the central dogma: An overview of mass spectrometry-based methods to investigate protein-metabolite interactions. Proteomics 2023, *n/a* (n/a), 2200533. DOI: 10.1002/pmic.202200533.

(4) Li, S.; Shui, W. Systematic mapping of protein–metabolite interactions with mass spectrometry-based techniques. Current Opinion in Biotechnology 2020, 64, 24–31. DOI: 10.1016/j.copbio.2019.09.002.

(5) Hicks, K. G.; Cluntun, A. A.; Schubert, H. L.; Hackett, S. R.; Berg, J. A.; Leonard, P. G.; Ajalla Aleixo, M. A.; Zhou, Y.; Bott, A. J.; Salvatore, S. R.;, et al. Protein-metabolite interactomics of carbohydrate metabolism reveal regulation of lactate dehydrogenase. Science 2023, 379 (6636), 996–1003. DOI: doi:10.1126/science.abm3452.

(6) Tian, Y.; Wan, N.; Zhang, H.; Shao, C.; Ding, M.; Bao, Q.; Hu, H.; Sun, H.; Liu, C.; Zhou, K.;, et al. Chemoproteomic mapping of the glycolytic targetome in cancer cells. Nature Chemical Biology 2023. DOI: 10.1038/s41589-023-01355-w.

(7) Venegas-Molina, J.; Molina-Hidalgo, F. J.; Clicque, E.; Goossens, A. Why and How to Dig into Plant Metabolite–Protein Interactions. Trends in Plant Science 2021, 26 (5), 472–483. DOI: 10.1016/j.tplants.2020.12.008.

(8) Diether, M.; Sauer, U. Towards detecting regulatory protein–metabolite interactions. Current Opinion in Microbiology 2017, 39, 16–23. DOI: 10.1016/j.mib.2017.07.006.

(9) Savitski, M. M.; Reinhard, F. B. M.; Franken, H.; Werner, T.; Savitski, M. F.; Eberhard, D.; Molina, D. M.; Jafari, R.; Dovega, R. B.; Klaeger, S.;, et al. Tracking cancer drugs in living cells by thermal profiling of the proteome. Science 2014, 346 (6205), 1255784. DOI: doi:10.1126/science.1255784.

(10) Meng, H.; Ma, R.; Fitzgerald, M. C. Chemical Denaturation and Protein Precipitation Approach for Discovery and Quantitation of Protein–Drug Interactions. Analytical Chemistry 2018, 90 (15), 9249–9255. DOI: 10.1021/acs.analchem.8b01772.

(11) Miyagi, M.; Tanaka, K.; Watanabe, S.; Kondo, J.; Kishimoto, T. Identifying Protein–Drug Interactions in Cell Lysates Using Histidine Hydrogen Deuterium Exchange. Analytical Chemistry 2021, 93 (45), 14985–14995. DOI: 10.1021/acs.analchem.1c02283.

(12) Zhang, X.; Wang, Q.; Li, Y.; Ruan, C.; Wang, S.; Hu, L.; Ye, M. Solvent-Induced Protein Precipitation for Drug Target Discovery on the Proteomic Scale. Analytical Chemistry 2020, 92 (1), 1363–1371. DOI: 10.1021/acs.analchem.9b04531.

(13) Reinhard, F. B. M.; Eberhard, D.; Werner, T.; Franken, H.; Childs, D.; Doce, C.; Savitski, M. F.; Huber, W.; Bantscheff, M.; Savitski, M. M.;, et al. Thermal proteome profiling monitors ligand interactions with cellular membrane proteins. Nature Methods 2015, 12 (12), 1129–1131. DOI: 10.1038/nmeth.3652.

(14) Feng, Y.; De Franceschi, G.; Kahraman, A.; Soste, M.; Melnik, A.; Boersema, P. J.; de Laureto, P. P.; Nikolaev, Y.; Oliveira, A. P.; Picotti, P. Global analysis of protein structural changes in complex proteomes. Nature Biotechnology 2014, 32 (10), 1036–1044. DOI: 10.1038/nbt.2999.

(15) Piazza, I.; Kochanowski, K.; Cappelletti, V.; Fuhrer, T.; Noor, E.; Sauer, U.; Picotti, P. A Map of Protein-Metabolite Interactions Reveals Principles of Chemical Communication. Cell 2018, 172 (1), 358–372.e323. DOI: 10.1016/j.cell.2017.12.006.

(16) Ma, R.; Meng, H.; Wiebelhaus, N.; Fitzgerald, M. C. Chemo-Selection Strategy for Limited Proteolysis Experiments on the Proteomic Scale. Analytical Chemistry 2018, 90 (23), 14039–14047. DOI: 10.1021/acs.analchem.8b04122.

(17) West, G. M.; Tucker, C. L.; Xu, T.; Park, S. K.; Han, X.; Yates, J. R.; Fitzgerald, M. C. Quantitative proteomics approach for identifying protein–drug interactions in complex mixtures using protein stability measurements. Proceedings of the National Academy of Sciences 2010, 107 (20), 9078–9082. DOI: doi:10.1073/pnas.1000148107.

(18) West, G. M.; Tang, L.; Fitzgerald, M. C. Thermodynamic Analysis of Protein Stability and Ligand Binding Using a Chemical Modification- and Mass Spectrometry-Based Strategy. Analytical Chemistry 2008, 80 (11), 4175–4185. DOI: 10.1021/ac702610a.

(19) Yan, W.; Wang, D.; Wan, N.; Wang, S.; Shao, C.; Zhang, H.; Zhao, Z.; Lu, W.; Tian, Y.; Ye, H.;, et al. Living Cell-Target Responsive Accessibility Profiling Reveals Silibinin Targeting ACSL4 for Combating Ferroptosis. Analytical Chemistry 2022, 94 (43), 14820–14826. DOI: 10.1021/acs.analchem.2c03515.

(20) Bennett, J. L.; Nguyen, G. T. H.; Donald, W. A. Protein–Small Molecule Interactions in Native Mass Spectrometry. Chemical Reviews 2022, 122 (8), 7327–7385. DOI: 10.1021/acs.chemrev.1c00293.

(21) Liu, P.; Zhang, J.; Ferguson, C. N.; Chen, H.; Loo, J. A. Measuring Protein–Ligand Interactions Using Liquid Sample Desorption Electrospray Ionization Mass Spectrometry. Analytical Chemistry 2013, 85 (24), 11966–11972. DOI: 10.1021/ac402906d.

(22) Cubrilovic, D.; Zenobi, R. Influence of Dimehylsulfoxide on Protein–Ligand Binding Affinities. Analytical Chemistry 2013, 85 (5), 2724–2730. DOI: 10.1021/ac303197p.

(23) Gavriilidou, A. F. M.; Gülbakan, B.; Zenobi, R. Influence of Ammonium Acetate Concentration on Receptor–Ligand Binding Affinities Measured by Native Nano ESI-MS: A Systematic Study. Analytical Chemistry 2015, 87 (20), 10378–10384. DOI: 10.1021/acs.analchem.5b02478.

(24) Maple, H. J.; Garlish, R. A.; Rigau-Roca, L.; Porter, J.; Whitcombe, I.; Prosser, C. E.; Kennedy, J.; Henry, A. J.; Taylor, R. J.; Crump, M. P.;, et al. Automated Protein–Ligand Interaction Screening by Mass Spectrometry. Journal of Medicinal Chemistry 2012, 55 (2), 837–851. DOI: 10.1021/jm201347k.

(25) Reher, R.; Aron, A. T.; Fajtová, P.; Stincone, P.; Wagner, B.; Pérez-Lorente, A. I.; Liu, C.; Shalom, I. Y. B.; Bittremieux, W.; Wang, M.;, et al. Native metabolomics identifies the rivulariapeptolide family of protease inhibitors. Nature Communications 2022, 13 (1), 4619. DOI: 10.1038/s41467-022-32016-6.

(26) Jecklin, M. C.; Touboul, D.; Bovet, C.; Wortmann, A.; Zenobi, R. Which electrospray-based ionization method best reflects protein-ligand interactions found in solution? A comparison of ESI, nanoESI, and ESSI for the determination of dissociation constants with mass spectrometry. Journal of the American Society for Mass Spectrometry 2008, 19 (3), 332–343. DOI: 10.1016/j.jasms.2007.11.007.

(27) Zheng, Q.; Tian, Y.; Ruan, X.; Chen, H.; Wu, X.; Xu, X.; Wang, G.; Hao, H.; Ye, H. Probing specific ligand-protein interactions by native-denatured exchange mass spectrometry. Anal Chim Acta 2018, 1036, 58–65. DOI: 10.1016/j.aca.2018.07.072.

(28) Nguyen, G. T. H.; Tran, T. N.; Podgorski, M. N.; Bell, S. G.; Supuran, C. T.; Donald, W. A. Nanoscale Ion Emitters in Native Mass Spectrometry for Measuring Ligand–Protein Binding Affinities. Acs Central Sci 2019, 5 (2), 308–318. DOI: 10.1021/acscentsci.8b00787.

(29) Susa, A. C.; Xia, Z.; Williams, E. R. Native Mass Spectrometry from Common Buffers with Salts That Mimic the Extracellular Environment. Angewandte Chemie International Edition 2017, 56 (27), 7912–7915. DOI: 10.1002/anie.201702330.

(30) Báez Bolivar, E. G.; Bui, D. T.; Kitova, E. N.; Han, L.; Zheng, R. B.; Luber, E. J.; Sayed, S. Y.; Mahal, L. K.; Klassen, J. S. Submicron Emitters Enable Reliable Quantification of Weak Protein–Glycan Interactions by ESI-MS. Analytical Chemistry 2021, 93 (9), 4231–4239. DOI: 10.1021/acs.analchem.0c05003.

(31) Yuill, E. M.; Sa, N.; Ray, S. J.; Hieftje, G. M.; Baker, L. A. Electrospray Ionization from Nanopipette Emitters with Tip Diameters of Less than 100 nm. Analytical Chemistry 2013, 85 (18), 8498–8502. DOI: 10.1021/ac402214g.

(32) Konermann, L.; Liu, Z.; Haidar, Y.; Willans, M. J.; Bainbridge, N. A. On the Chemistry of Aqueous Ammonium Acetate Droplets during Native Electrospray Ionization Mass Spectrometry. Analytical Chemistry 2023, 95 (37), 13957–13966. DOI: 10.1021/acs.analchem.3c02546.

(33) Davis, B. T. V.; Velyvis, A.; Vahidi, S. Fluorinated Ethylamines as Electrospray-Compatible Neutral pH Buffers for Native Mass Spectrometry. Analytical Chemistry 2023, 95 (48), 17525–17532. DOI: 10.1021/acs.analchem.3c02640.

(34) Hedges, J. B.; Vahidi, S.; Yue, X.; Konermann, L. Effects of Ammonium Bicarbonate on the Electrospray Mass Spectra of Proteins: Evidence for Bubble-Induced Unfolding. Analytical Chemistry 2013, 85 (13), 6469–6476. DOI: 10.1021/ac401020s.

(35) Zhang, H.; Lu, H.; Chingin, K.; Chen, H. Stabilization of Proteins and Noncovalent Protein Complexes during Electrospray Ionization by Amino Acid Additives. Analytical Chemistry 2015, 87 (14), 7433–7438. DOI: 10.1021/acs.analchem.5b01643.

(36) Townsend, J. A.; Keener, J. E.; Miller, Z. M.; Prell, J. S.; Marty, M. T. Imidazole Derivatives Improve Charge Reduction and Stabilization for Native Mass Spectrometry. Analytical Chemistry 2019, 91 (22), 14765–14772. DOI: 10.1021/acs.analchem.9b04263.

(37) Sun, J.; Kitova, E. N.; Wang, W.; Klassen, J. S. Method for Distinguishing Specific from Nonspecific Protein−Ligand Complexes in Nanoelectrospray Ionization Mass Spectrometry. Analytical Chemistry 2006, 78 (9), 3010–3018. DOI: 10.1021/ac0522005.

(38) Sun, N.; Soya, N.; Kitova, E. N.; Klassen, J. S. Nonspecific Interactions Between Proteins and Charged Biomolecules in Electrospray Ionization Mass Spectrometry. Journal of the American Society for Mass Spectrometry 2010, 21 (3), 472–481. DOI: 10.1016/j.jasms.2009.12.002.

(39) Fernandez de la Mora, J. Electrospray ionization of large multiply charged species proceeds via Dole’s charged residue mechanism. Anal Chim Acta 2000, 406 (1), 93–104. DOI: 10.1016/S0003-2670(99)00601-7.

(40) Konermann, L.; Ahadi, E.; Rodriguez, A. D.; Vahidi, S. Unraveling the Mechanism of Electrospray Ionization. Analytical Chemistry 2013, 85 (1), 2–9. DOI: 10.1021/ac302789c.

(41) Kebarle, P.; Verkerk, U. H. Electrospray: From ions in solution to ions in the gas phase, what we know now. Mass Spectrometry Reviews 2009, 28 (6), 898–917. DOI: 10.1002/mas.20247.

(42) Van Berkel, G. J.; Kertesz, V. An open port sampling interface for liquid introduction atmospheric pressure ionization mass spectrometry. Rapid Communications in Mass Spectrometry 2015, 29 (19), 1749–1756. DOI: 10.1002/rcm.7274.

(43) Looby, N. T.; Tascon, M.; Acquaro, V. R.; Reyes-Garcés, N.; Vasiljevic, T.; Gomez-Rios, G. A.; Wąsowicz, M.; Pawliszyn, J. Solid phase microextraction coupled to mass spectrometry via a microfluidic open interface for rapid therapeutic drug monitoring. Analyst 2019, 144 (12), 3721–3728, 10.1039/C9AN00041K. DOI: 10.1039/C9AN00041K.

(44) Tascon, M.; Singh, V.; Huq, M.; Pawliszyn, J. Direct Coupling of Dispersive Extractions with Magnetic Particles to Mass Spectrometry via Microfluidic Open Interface. Analytical Chemistry 2019, 91 (7), 4762–4770. DOI: 10.1021/acs.analchem.9b00308.

(45) Sosnowski, P.; Marin, V.; Tian, X.; Hopfgartner, G. Analysis of illicit pills and drugs of abuse in urine samples using a 3D-printed open port probe hyphenated with differential mobility spectrometry-mass spectrometry. Analyst 2022, 147 (19), 4318–4325, 10.1039/D2AN00925K. DOI: 10.1039/D2AN00925K.

(46) Sosnowski, P.; Hopfgartner, G. Application of 3D printed tools for customized open port probe-electrospray mass spectrometry. Talanta 2020, 215, 120894. DOI: 10.1016/j.talanta.2020.120894.

(47) Grandori, R. Origin of the conformation dependence of protein charge-state distributions in electrospray ionization mass spectrometry. Journal of Mass Spectrometry 2003, 38 (1), 11–15. DOI: 10.1002/jms.390.

(48) Šamalikova, M.; Grandori, R. Role of opposite charges in protein electrospray ionization mass spectrometry. Journal of Mass Spectrometry 2003, 38 (9), 941–947. DOI: 10.1002/jms.507.

(49) Testa, L.; Brocca, S.; Grandori, R. Charge-Surface Correlation in Electrospray Ionization of Folded and Unfolded Proteins. Analytical Chemistry 2011, 83 (17), 6459–6463. DOI: 10.1021/ac201740z.

(50) Liigand, J.; de Vries, R.; Cuyckens, F. Optimization of flow splitting and make-up flow conditions in liquid chromatography/electrospray ionization mass spectrometry. Rapid Communications in Mass Spectrometry 2019, 33 (3), 314–322. DOI: 10.1002/rcm.8352.

(51) Kostiainen, R.; Kauppila, T. J. Effect of eluent on the ionization process in liquid chromatography–mass spectrometry. Journal of Chromatography A 2009, 1216 (4), 685–699. DOI: 10.1016/j.chroma.2008.08.095.

(52) Zheng, Q.; Ruan, X.; Tian, Y.; Hu, J.; Wan, N.; Lu, W.; Xu, X.; Wang, G.; Hao, H.; Ye, H. Ligand–protein target screening from cell matrices using reactive desorption electrospray ionization-mass spectrometry via a native-denatured exchange approach. Analyst 2019, 144 (2), 512–520, 10.1039/C8AN01708E. DOI: 10.1039/C8AN01708E.

(53) Daniel, J. M.; Friess, S. D.; Rajagopalan, S.; Wendt, S.; Zenobi, R. Quantitative determination of noncovalent binding interactions using soft ionization mass spectrometry. International Journal of Mass Spectrometry 2002, 216 (1), 1–27. DOI: 10.1016/S1387-3806(02)00585-7.

(54) Barbiroli, A.; Beringhelli, T.; Bonomi, F.; Donghi, D.; Ferranti, P.; Galliano, M.; Iametti, S.; Maggioni, D.; Rasmussen, P.; Scanu, S.;, et al. Bovine β-lactoglobulin acts as an acid-resistant drug carrier by exploiting its diverse binding regions. Biological Chemistry 2010, 391 (1), 21–32. DOI: doi:10.1515/bc.2010.008 (acccessed 2023-12-05).

(55) Zhang, S.; Van Pelt, C. K.; Wilson, D. B. Quantitative Determination of Noncovalent Binding Interactions Using Automated Nanoelectrospray Mass Spectrometry. Analytical Chemistry 2003, 75 (13), 3010–3018. DOI: 10.1021/ac034089d.

