## Supplementary Infos for "Differentiating specific and non-specific protein-metabolite interactions using gradient open port probe electrospray ionization mass spectrometry"

**Table of contents**

Figure S1. Ranking the affinity of CTP, CDP, and CMP to RNase A

Figure S2. Measuring the Kd of CTP and CDP to RNase A by titration assays.

Figure S3. The BPC of RNase A for the OPP gradient from 0 to 90% MeOH.

Figure S4. Reproducibility of the gradient-OPP-ESI-MS approach.

Figure S5. Triplicates analyses of the interaction between CDP and RNase A under the native conditions.

Figure S6. Triplicates analyses of the interaction between CDP and RNase A under gradient.

Figure S7. Intensity ratio of RNase A and RNase A-CDP complex around 70% MeOH.

Figure S8. Differentiations between the specific PMIs and non-specific bindings with lysozyme.

Figure S9. Differentiations between the specific PMIs and non-specific bindings with beta-lactoglobulin.

Figure S10. Comparation between the specific PMIs (lysozyme-NTAC) and nonspecific adhesions (lysozyme-CTP) in CSD over the OPP gradient.

Figure S11. Comparation between the specific PMIs (beta-lactoglobulin-fluvastatin) and nonspecific adhesions (beta-lactoglobulin-CTP) in CSD over the OPP gradient.

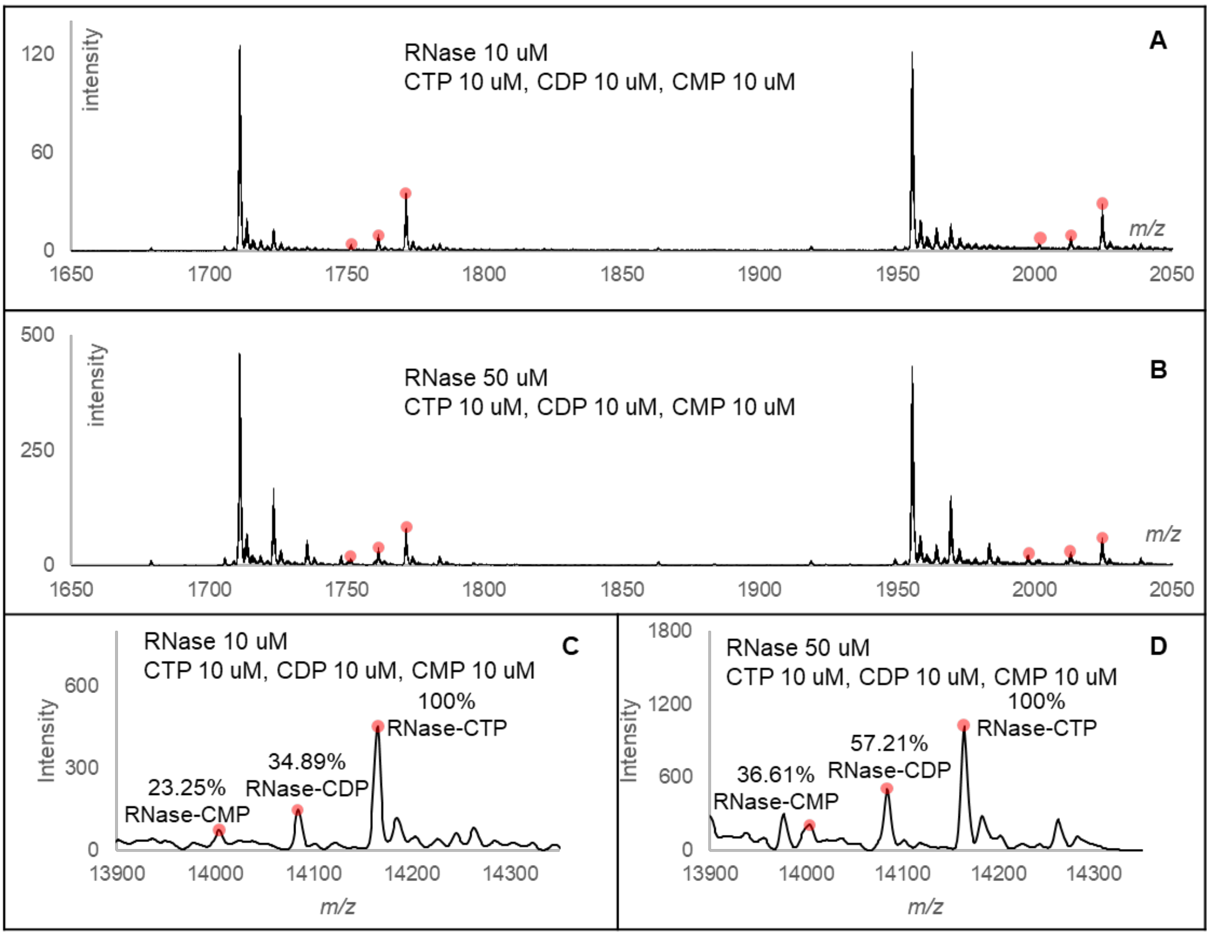


**Figure S1**. Ranking the affinity of CTP, CDP, and CMP to RNase A.

**A**. the raw spectrum of competitive experiments, in which three ligands have the same concentration of 10 µM and the concentration of RNase A is 10 µM that is lower than the total concentration of ligands.

**B**. the raw spectrum of noncompetitive experiments, in which three ligands have the same concentration of 10 µM and the concentration of RNase A is 50 µM that is higher than the total concentration of ligands.

**C**. the reconstructed spectrum of competitive experiments after deconvolution.

**D**. the reconstructed spectrum of noncompetitive experiments after deconvolution.

| C. (RNase A, uM) | C. (CTP, uM) | R ([PL]/[P]) |
|---|---|---|
| 5 | 0.5 | 0.0434 |
| 5 | 1 | 0.0628 |
| 5 | 2 | 0.0823 |
| 5 | 5 | 0.2776 |
| 5 | 10 | 0.5001 |
| 5 | 15 | 0.7850 |
| 5 | 20 | 0.7904 |
| 5 | 35 | 1.2023 |
| 5 | 50 | 2.2550 |

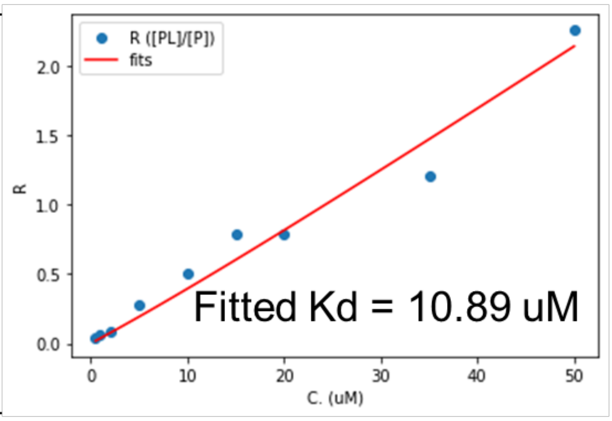


| C. (RNase A, uM) | C. (CDP, uM) | R ([PL]/[P]) |
|---|---|---|
| 5 | 2 | 0.0280 |
| 5 | 5 | 0.0911 |
| 5 | 10 | 0.1233 |
| 5 | 15 | 0.1926 |
| 5 | 20 | 0.2866 |
| 5 | 35 | 0.5130 |
| 5 | 50 | 0.5463 |

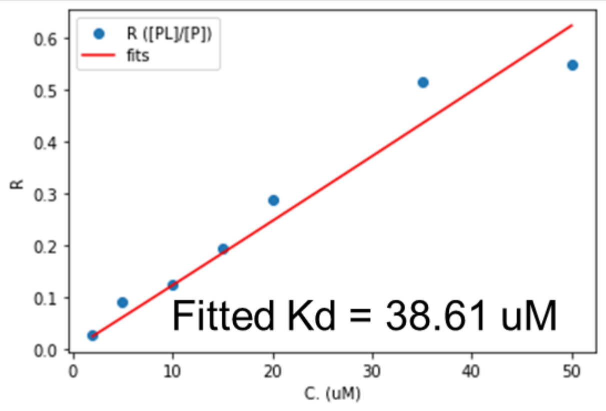


Measured R values versus the concentration of ligands (CTP/CDP) were fitted by the nonlinear least squares method to fit the function below.

$$R = \frac{-(K_d - 0.5 \cdot [CXP] + 0.5 \cdot [RNaseA]) + \sqrt{(K_d - 0.5 \cdot [CXP] + 0.5[RNaseA])^2 + 2 \cdot K_d \cdot [CXP]}}{2 \cdot K_d}$$

**Figure S2**. Measuring the Kd of CTP and CDP to RNase A by titration assays. The concentration of RNase A is constant at 5 µM. The ligands of CTP and CDP cover concentrations range from 0.5 µM to 50 µM. For each ligand concentration, the *R* is calculated by dividing the peak area of complex [PL] with the peak area of free protein [P]. The generated *R* values are plotted versus the concentrations of ligands. With the equation above, the points are fitted by the nonlinear least squares methods. The measured Kd is 10.89 µM for CTP and 38.61 µM for CDP.

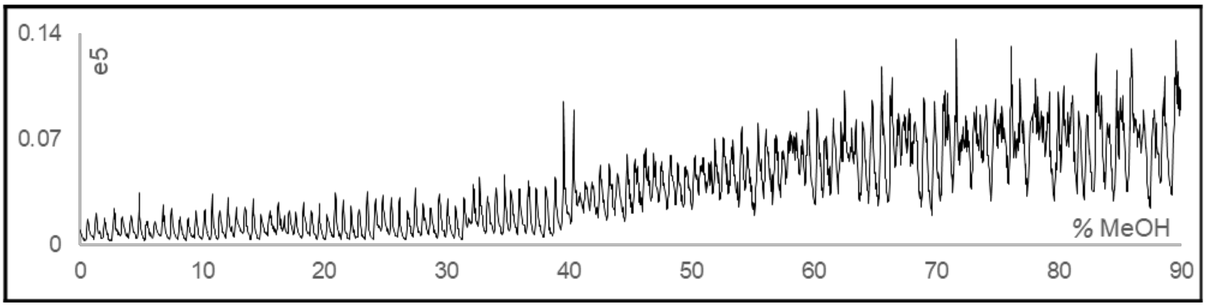


**Figure S3**. The BPC of RNase A for the OPP gradient from 0 to 90% MeOH.

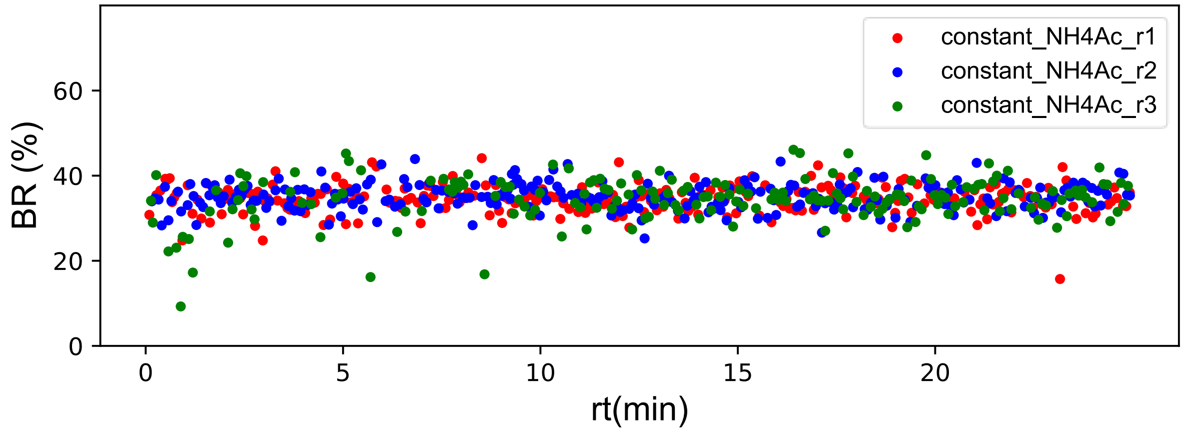


**Figure S4**. Reproducibility of the gradient-OPP-ESI-MS approach. Triplicate analyses of CDP interacting with RNase A at constant 10 mM $NH_4Ac$ OPP mobile phase. Binding ratio, BR is the ratio [PL]/([PL] + [P]).

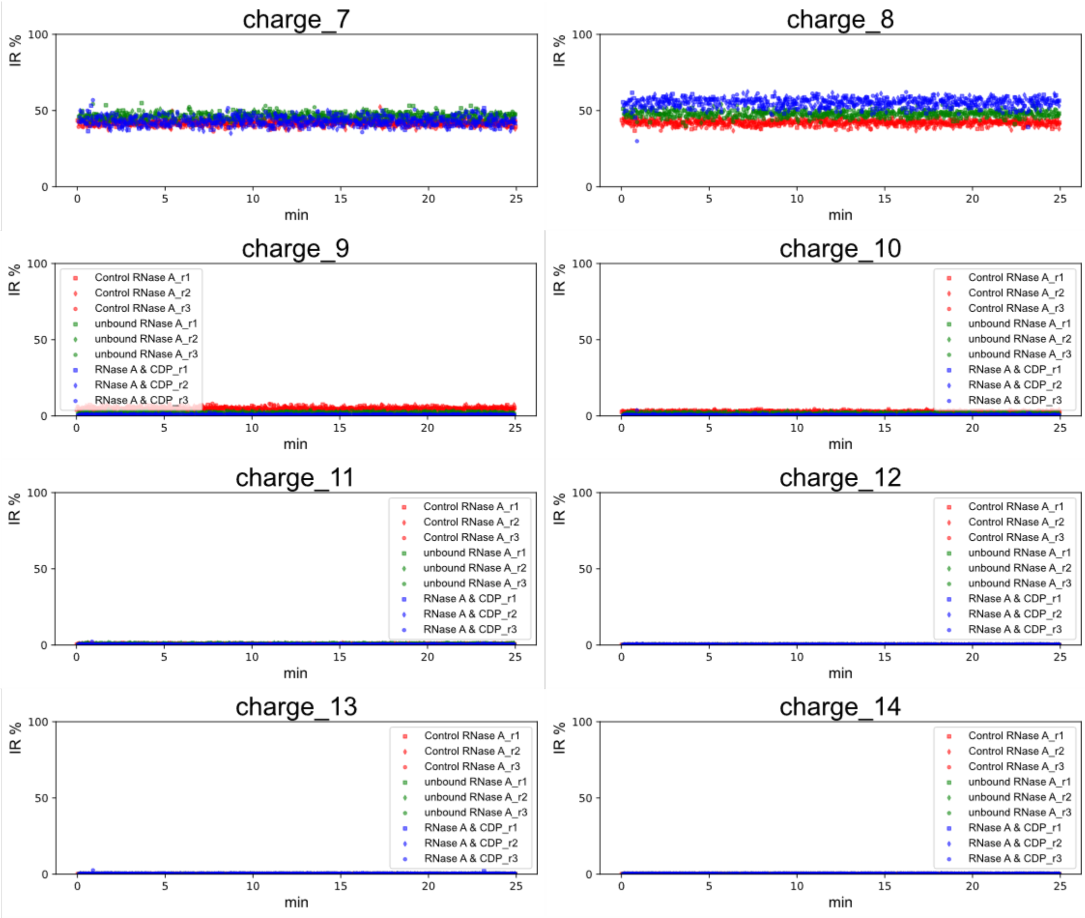


**Figure S5**. Triplicates analyses of the interaction between CDP and RNase A under the native conditions. The intensity ratios of all charges remain unchanged. There are two samples, the control sample containing RNase A without CDP and the experimental sample containing both RNase A and CDP. The RNase A in the control sample was labeled as “control-RNase A” and the free RNase A in the experimental sample was labeled as “unbound-RNase A”.

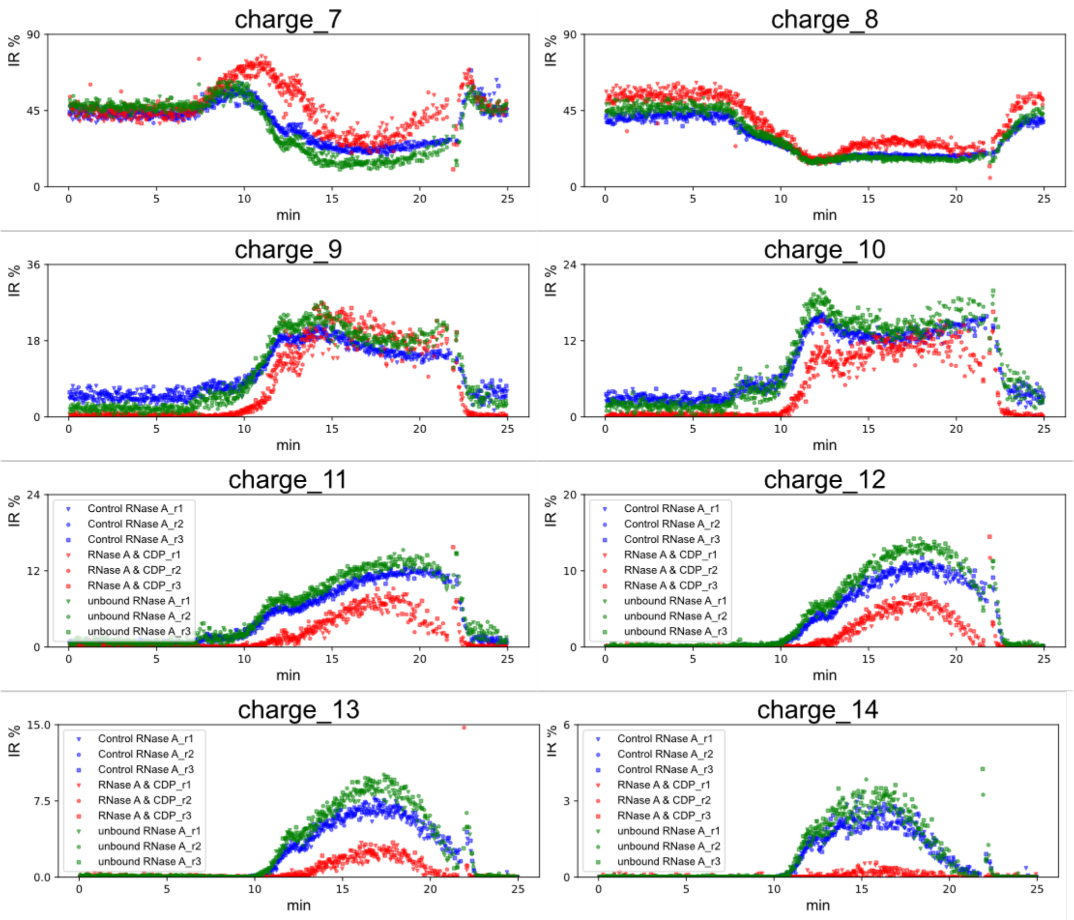


**Figure S6**. Triplicates analyses of the interaction between CDP and RNase A under a gradient, that consist of 0-5 min, constant at 100% 10 mM NH4Ac, 5-10 min, change from 100% 10 mM $NH_4Ac$ to 100% $H_2O$ containing 0.1% formic acid, 10-20 min, change from 0 to 90% MeOH. 20-25 min, equilibrate at 100% 10 mM $NH_4Ac$. The spikes around 21 min may be caused by the sudden change from 90% MeOH to 100% 10 mM $NH_4Ac$.

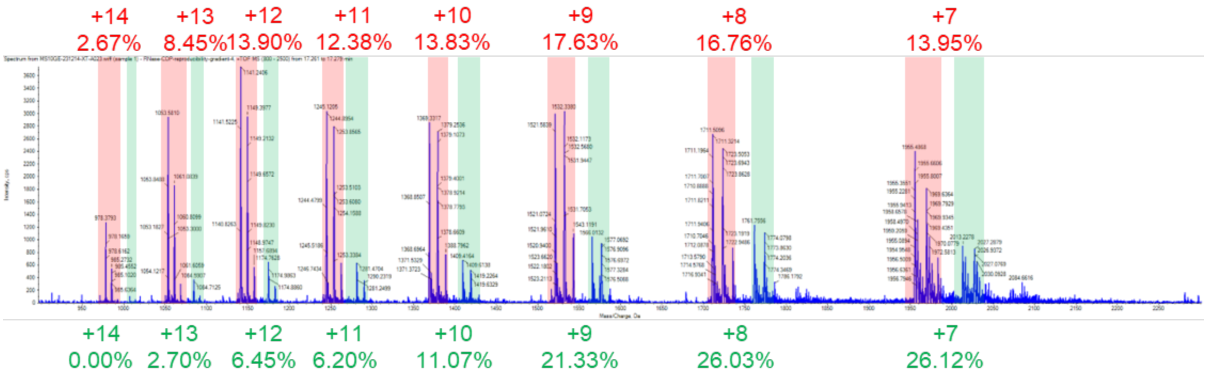


| | +14 | +13 | +12 | +11 | +10 | +9 | +8 | +7 |
|---|---|---|---|---|---|---|---|---|
| **RNase A** | 2.67 | 8.45 | 13.90 | 12.38 | 13.83 | 17.63 | 16.76 | 13.95 |
| **RNase A-CDP** | 0.00 | 2.70 | 6.45 | 6.20 | 11.07 | 21.33 | 26.03 | 26.12 |

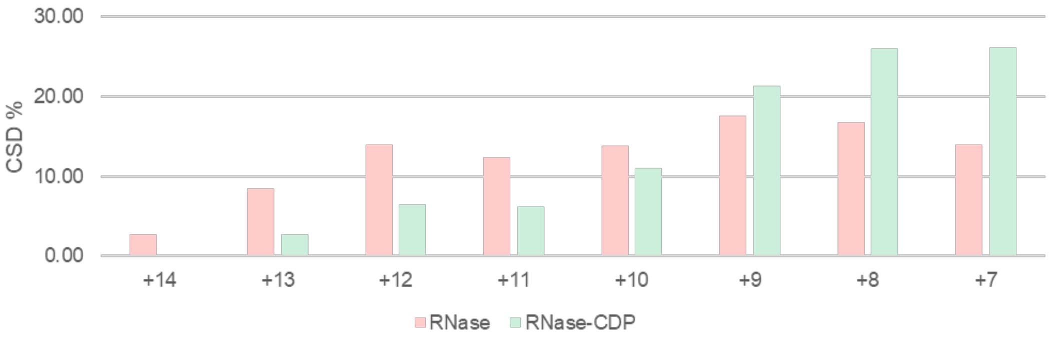


**Figure S7**. The MS spectrum was taken at around 70% MeOH. Both RNase A-CDP complex and unbound-RNase existed in the same spectrum. Intensity ratios of all charge states for both were calculated and plotted in bar chat. The CSD shapes are obviously different.

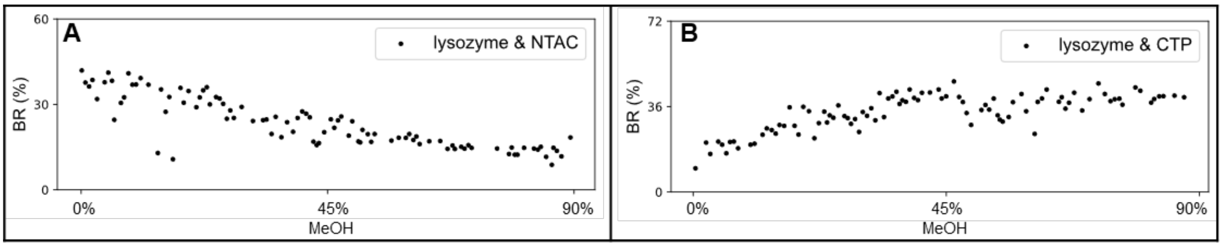


**Figure S8.** Differentiations between the specific PMIs and non-specific bindings. **A**. binding ratios of lysozyme and NTAC; **B**. binding ratios of lysozyme and CTP.

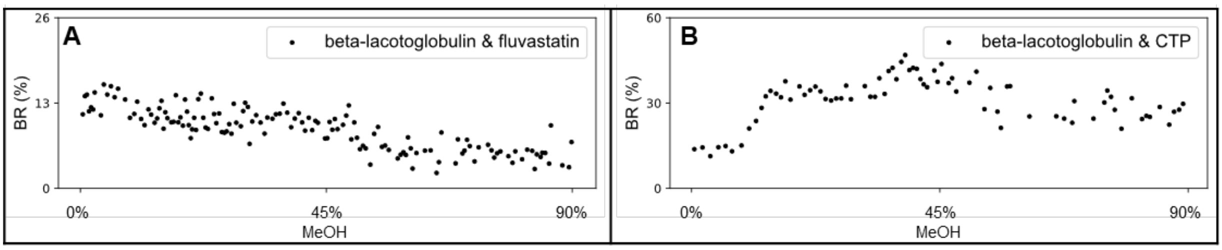


**Figure S9.** Differentiations between the specific PMIs and non-specific bindings. **A**. binding ratios of beta-lactoglobulin and fluvastatin; **B**. binding ratios of beta-lactoglobulin and CTP.

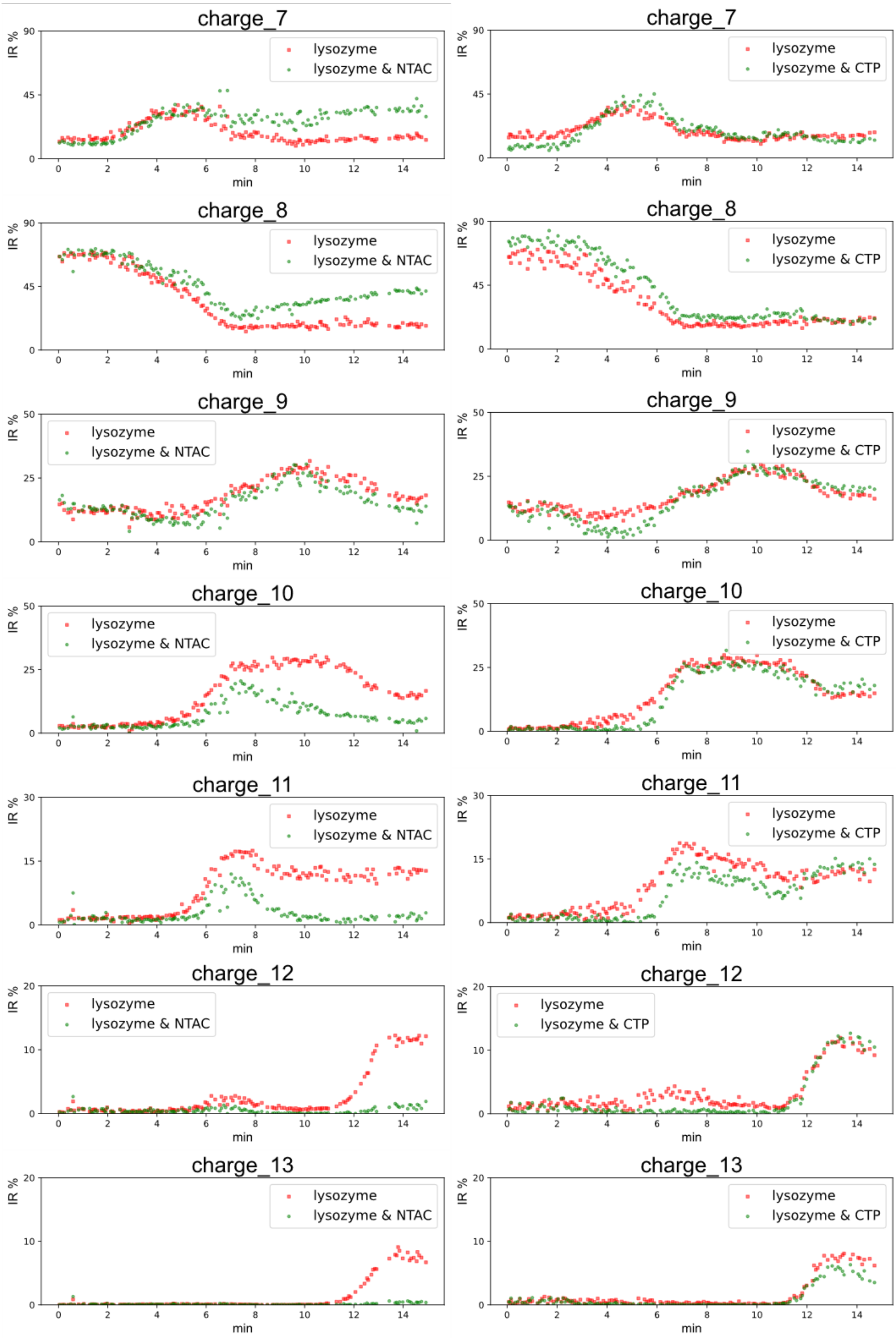


**Figure S10**. Comparation between the specific PMIs (lysozyme-NTAC) and nonspecific adhesions (lysozyme-CTP) in CSD over the OPP gradient.

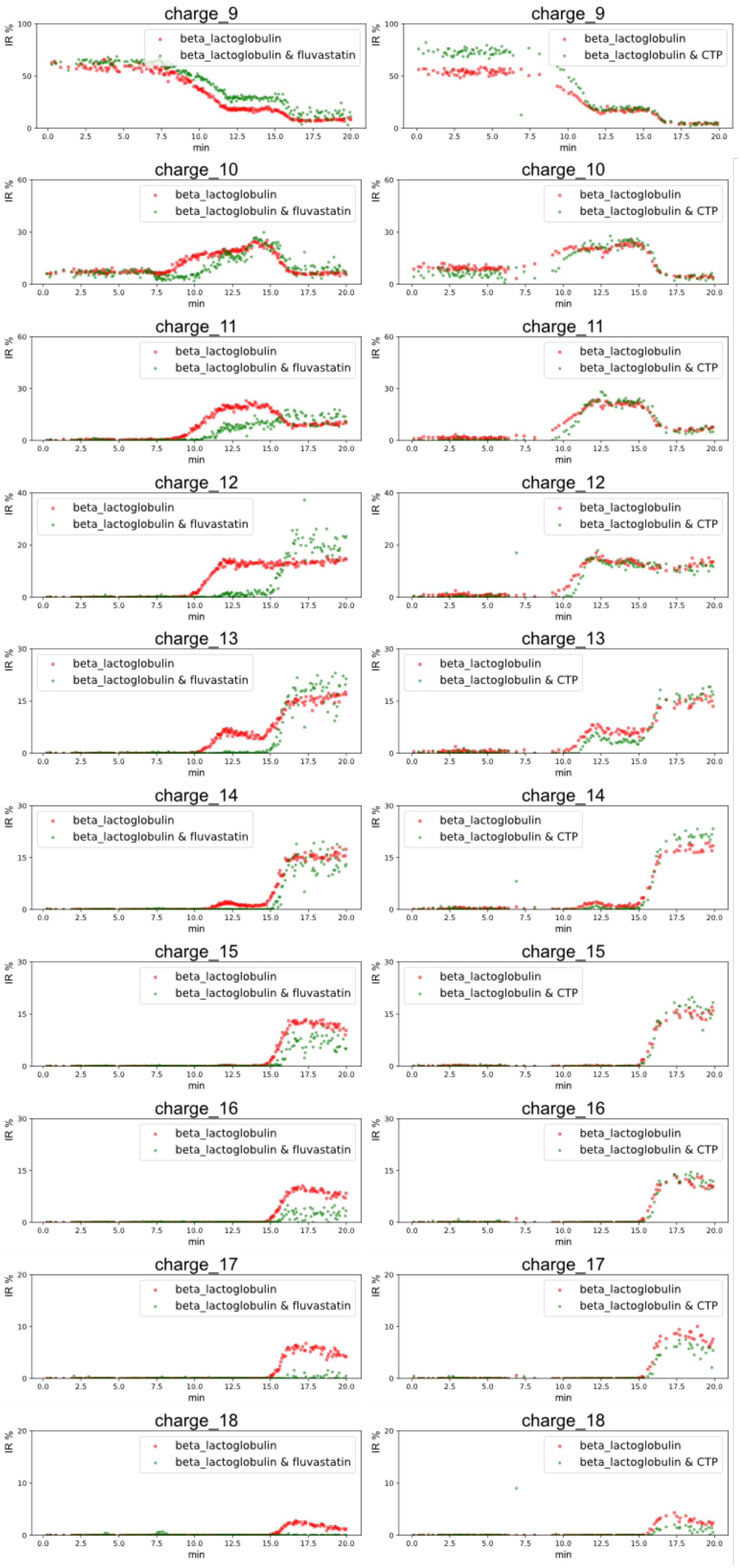


**Figure S11**. Comparation between the specific PMIs (beta-lactoglobulin-fluvastatin) and nonspecific adhesions (beta-lactoglobulin-CTP) in CSD over the OPP gradient.
